# Identifying inhibitors of epithelial-mesenchymal plasticity using a network topology based approach

**DOI:** 10.1101/854307

**Authors:** Kishore Hari, Burhanuddin Sabuwala, Balaram Vishnu Subramani, Caterina La Porta, Stefano Zapperi, Francesc Font-Clos, Mohit Kumar Jolly

## Abstract

Metastasis is the cause of over 90% of cancer-related deaths. Cancer cells undergoing metastasis can switch dynamically between different phenotypes, enabling them to adapt to harsh challenges such as overcoming anoikis and evading immune response. This ability, known as phenotypic plasticity, is crucial for the survival of cancer cells during metastasis, as well as acquiring therapy resistance. Various biochemical networks have been identified to contribute to phenotypic plasticity, but how plasticity emerges from the dynamics of these networks remains elusive. Here, we investigated the dynamics of various regulatory networks implicated in Epithelial-Mesenchymal Plasticity (EMP) − an important arm of phenotypic plasticity − through two different mathematical modelling frameworks: a discrete, parameter-independent framework (Boolean) and a continuous, parameter-agnostic modelling framework (RACIPE). Results from either framework in terms of phenotypic distributions obtained from a given EMP network are qualitatively similar and suggest that these networks are multi-stable and can give rise to phenotypic plasticity. Neither method requires specific kinetic parameters, thus our results emphasize that EMP can emerge through these networks over a wide range of parameter sets, elucidating the importance of network topology in enabling phenotypic plasticity. Furthermore, we show that the ability to exhibit phenotypic plasticity correlates positively with the number of positive feedback loops in a given network. These results pave a way towards an unorthodox network topology-based approach to identify crucial links in a given EMP network that can reduce phenotypic plasticity and possibly inhibit metastasis - by reducing the number of positive feedback loops.

## 1. Introduction

Metastasis, therapy resistance, and tumor relapse remain unsolved clinical challenges and major causes of cancer mortality [1]. During metastasis, cells navigate many bottlenecks: local invasion, intravasation, survival in circulation in matrix-deprived conditions, extravasation, and eventually colonization of the distant organ. Only a few (< 0.02%) cells survive this cascade of events and are capable of initiating metastasis. Recent studies have identified phenotypic plasticity − the ability of cells to reversibly switch phenotypes in response to their ever-changing environmental conditions − as a hallmark of cancer metastasis [2]. Similarly, phenotypic plasticity enables a small proportion of cancers cells to transiently acquire an adaptive drug-refractory phenotype which may contribute to tumor relapse [3]. Therefore, identifying the mechanisms of phenotypic plasticity is essential for any major breakthroughs in cancer treatment.

Phenotypic plasticity is considered to be an adaptation strategy to survive in variable environmental conditions [4]. Recently, the contributions of phenotypic plasticity in driving metastasis and therapy resistance have been realized more prominently, especially due to the lack of any unique mutational signature being identified for cancer metastasis so far [2], and the frequent emergence of resistance against targeted therapy [3]. Phenotypic plasticity, referred to as “the architect who never sleeps” [5], can have various dimensions – metabolic plasticity [6], epithelial-mesenchymal plasticity [7], plasticity between cancer stem cell (CSC) and a non-CSC state [8, 9, 10], and plasticity between drug-sensitive and drug-resistant/tolerant state [11] among others. Cancer cells continually exploit phenotypic plasticity to adapt to their ever-changing environment by maximizing their fitness during cancer progression, metastasis, therapy resistance and eventually tumor relapse [12]. Moreover, phenotypic plasticity can amplify the non-genetic heterogeneity within tumors, thus increasing the number of ‘exit options’ for cells in response to drugs, thereby increasing the versatility of the tumor cell population [13]. Thus, targeting phenotypic plasticity provides a unique opportunity to both curb cancer metastasis and to improve the efficiency of existing therapeutic strategies.

A canonical example of phenotypic plasticity is Epithelial-Mesenchymal plasticity (EMP), which involves partial and/or complete Epithelial-Mesenchymal Transition (EMT) and/or Mesenchymal-Epithelial Transition (MET). EMT and MET are embryonic developmental programs which are often adopted by cancer cells during metastasis [14]. EMT can fuel the dissemination of stationary cancer cells through reduced cell-cell adhesion and apico-basal polarity and increased migration and invasion. Conversely, MET may enable the disseminated cells that exit the bloodstream to survive in a different environment by regaining cell-cell adhesion and proliferation to eventually colonize that organ. EMT and MET were earlier thought of as binary processes (i.e. switching between epithelial and mesenchymal phenotypes), but recent *in vitro, in vivo* and *in silico* evidence strongly suggests that they are not ‘all-or-none’ processes, and cells can stably reside in one or more hybrid E/M phenotypes, thus enabling various manifestations of EMP [14]. EMP can not only drive metastasis, but also influence the resistance to various chemotherapeutic drugs and targeted therapies. It can also alter the tumor microenvironment to be immunosuppressive, eventually leading to overall poor survival of patients across cancer types [15, 16]. Therefore, preventing the ability of cells to reversibly switch among these epithelial (E), mesenchymal (M), and hybrid E/M (H) phenotypes can have a significant clinical impact.

Most current preclinical and clinical efforts attempt to restrict EMP in only one direction – EMT or MET. Such efforts are likely to facilitate the transition in the opposite direction (i.e. MET or EMT respectively) and can possibly increase the frequency of hybrid E/M phenotypes which can be the ‘fittest’ for metastasis [17]. Hence, these interventions may increase the metastatic load instead, depending on the phenotypic distribution of disseminated cells. Blocking EMP in both directions can overcome these potential side effects. Moreover, restricting EMP bidirectionally can also limit the ability of a clonal population to generate and/or maintain non-genetic heterogeneity [18]. Non-genetic heterogeneity can enable ‘bet-hedging’ during the evolution of drug resistance [19], as well as the co-existence of phenotypically distinct subpopulations of epithelial and mesenchymal cells that can communicate and cooperate among themselves to aggravate metastasis [20, 21]. Thus, blocking EMP bidirectionally, or in other words “fixing cells at a given position on the epithelial-–mesenchymal axis to prevent access to the range of states that might be required to facilitate different stages of the metastatic cascade” [22], is likely to blunt the metastatic and drug-resistance potential of cancer cells much stronger than restricting only EMT or only MET.

Identifying ways to inhibit EMP requires a detailed mechanistic understanding of its dynamics. Experimentally, EMP can be tracked through recent advances such as live-cell imaging reporter constructs or single-cell RNA-seq and/or mass/flow cytometry [13]. Another approach to characterize the dynamics of EMP is by developing mechanism-based mathematical models of networks that have experimentally been identified to regulate EMT and/or MET. Many such mathematical models have helped gain useful insights into the dynamics of EMP and have driven the experiments to decode a) how cells attain one or more hybrid E/M phenotype, b) how reversible is EMP in both directions (E to M vs. M to E) and c) whether cells take same or different paths en route EMT or MET [23]. However, none of these models investigated the possibility of identifying mechanisms to block EMP bidirectionally.

Here, we investigate the dynamics of various regulatory networks implicated in EMP and identify robust network topology based design principles of EMP, using two different but complementary modeling formalisms – a discrete, parameter-independent method: asynchronous Boolean [24] and a continuous, parameter-agnostic method: RACIPE (Random Circuit Perturbation) [25]. Our results show that the phenotypic distributions that can be obtained through an EMP network depend majorly on network topology but are largely independent of specific kinetic parameters for each link in the network. We also pinpoint a set of network perturbations that can reduce EMP, and observe a unifying theme amongst them: a reduced number of total positive feedback loops embedded within an EMP network led to curtailed EMP. Therefore, our approach unravels the common operating principles of various EMP regulatory networks and offers a systematic framework to identify network perturbations to restrict EMP based on this network topology-based dynamical trait.

## 2. Results

### 2.1. RACIPE and Boolean models have similar phenotypic distributions for EMP networks

Boolean frameworks lack kinetic parameters and treat a gene to be discretely ON (1) or OFF (0), thus focusing on a coarse-grained view of how various interactions in a network can give rise to the repertoire of dynamical behaviours [26]. RACIPE, on the other hand, generates an ensemble of continuous mathematical models (sets of coupled ordinary differential equations) with randomly chosen kinetic parameters for a given network topology and clusters the steady state solutions to identify the robust dynamical features of a given network. In other words, while a Boolean framework is parameter-independent, RACIPE can be thought of as a parameter-agnostic one.

Therefore, the similarities in dynamical traits of a network simulated via Boolean and RACIPE frameworks can unravel the extent to which the network topology drives the network dynamics, without much reliance on the specific choice of kinetic parameters. Across various EMP networks and the perturbations made in those, we have compared the outputs of these two modeling frameworks in terms of phenotypic distributions, and in ranking the effect of various perturbations in diminishing EMP (**Fig 1A**).

**Figure 1:**
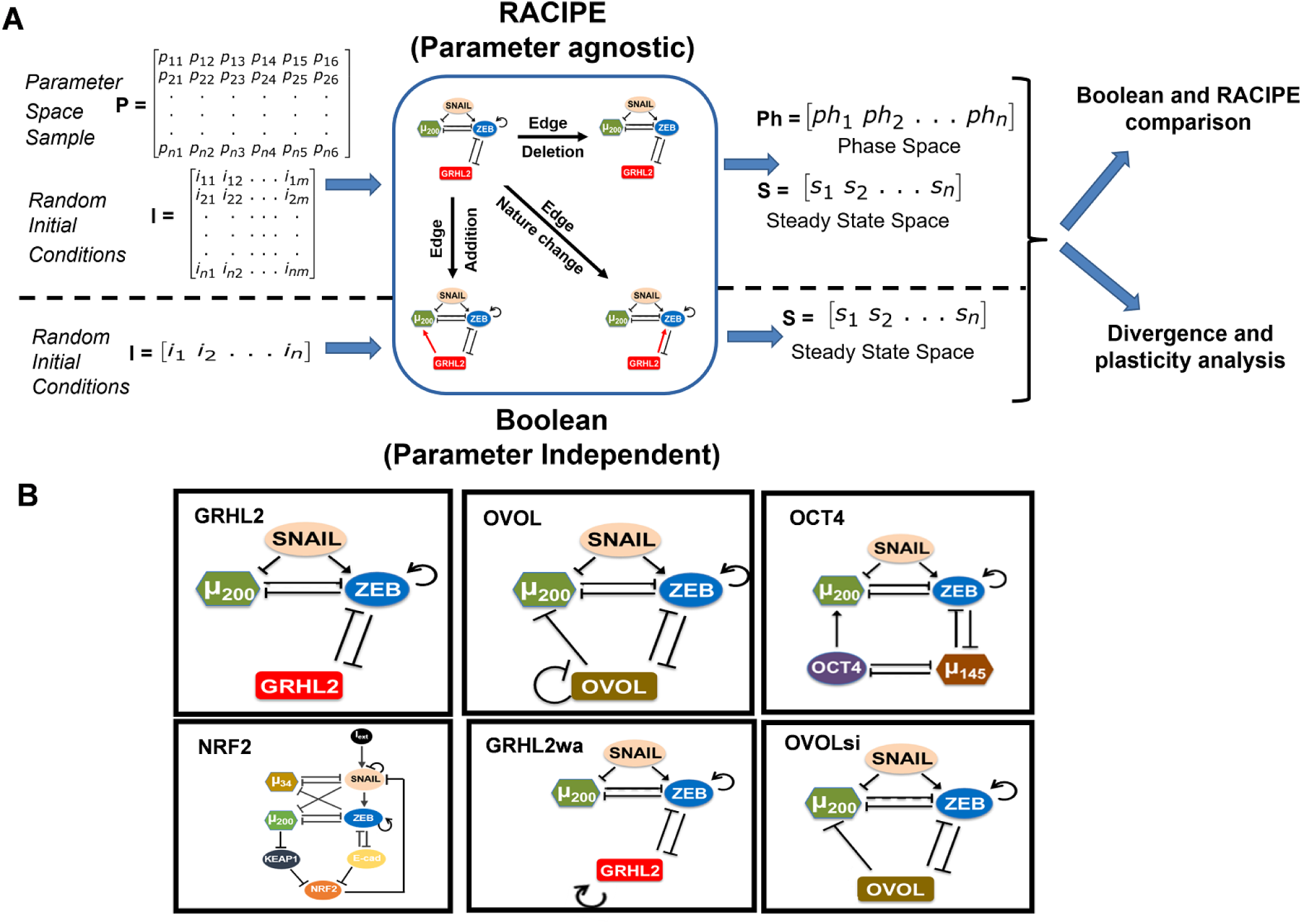
Dynamical approaches to investigate EMP. A. Schematic of network analysis strategy. The top part of the figure depicts RACIPE methodology of sampling random parameter sets (matrix P) and for each parameter set (each row in the matrix), randomly sampling multiple initial conditions to obtain steady states space and the phase space, i.e, information about monostable vs multi-stable parameter regions or phases. The bottom part of the figure depicts Boolean simulation method where for a given network, multiple initial conditions are randomly chosen and steady state space is obtained through asynchronous update. The center square depicts various possible topological perturbations done to the networks. B. EMP networks analysed in the study, namely, GRHL2 (top-left, 4 nodes, 7 edges), OVOL (top-center, 4 nodes, 9 edges), OCT4 [27] (top-right, 5 nodes, 10 edges), NRF2 [28] (bottom-left, 8 nodes, 16 edges), GRHL2wa (bottom-center, 4 nodes, 8 edges) and OVOLsi [27] (bottom-right, 4 nodes, 8 edges).

We have investigated 6 different networks reported in EMP literature; these networks vary from 3 nodes to 8 nodes and 7 edges to 16 edges (**Fig 1B**). First, we calculated the phenotypic distributions (i.e. stable steady state frequency distributions) obtained via RACIPE and Boolean models. To facilitate the comparison of Boolean and RACIPE models, we have discretized the output of RACIPE (as described in ‘Methods’ section). First, we determined the sample size of parameter sets to be chosen for RACIPE, and the number of initial conditions for Boolean models, using a quantitative convergence analysis. N=10,000 was chosen as the optimal number of parameter sets for RACIPE, and as the optimal number of initial conditions for Boolean analysis, based on observed standard deviation in steady state distributions obtained from RACIPE and Boolean models (**Fig 2A, S1A**).

For the miR-200/ZEB/ SNAIL/GRHL2 network (hereafter called as ‘GRHL2 network’; 4 nodes, 7 edges), a maximum of 2^3^ = 8 stable steady states are possible (value of each node = 0 or 1; SNAIL is an input to the circuit), in discretized RACIPE and Boolean framework. From Boolean analysis, we obtained four stable states for this network across different numbers of initial conditions chosen. Two out of these four states were more prominent − (ZEB=0, miR-200=1, GRHL2=1) and (ZEB=1, miR-200=0, GRHL2=0) − than the others (**Fig 2A,i**). These two states can be construed as epithelial (high miR-200 and GRHL2, low ZEB) and mesenchymal (low miR-200 and GRHL2, high ZEB) phenotypes as observed experimentally [29, 30]. Discretized analysis of RACIPE results also identifies these four stable steady states with similar relative frequency as seen in the case of Boolean model, and 3 other states with relatively less frequencies (**Fig 2A,ii**). Put together, these results suggest that epithelial and mesenchymal phenotypes are the two most commonly expected phenotypes from the dynamics of GRHL2 network.

**Figure 2:**
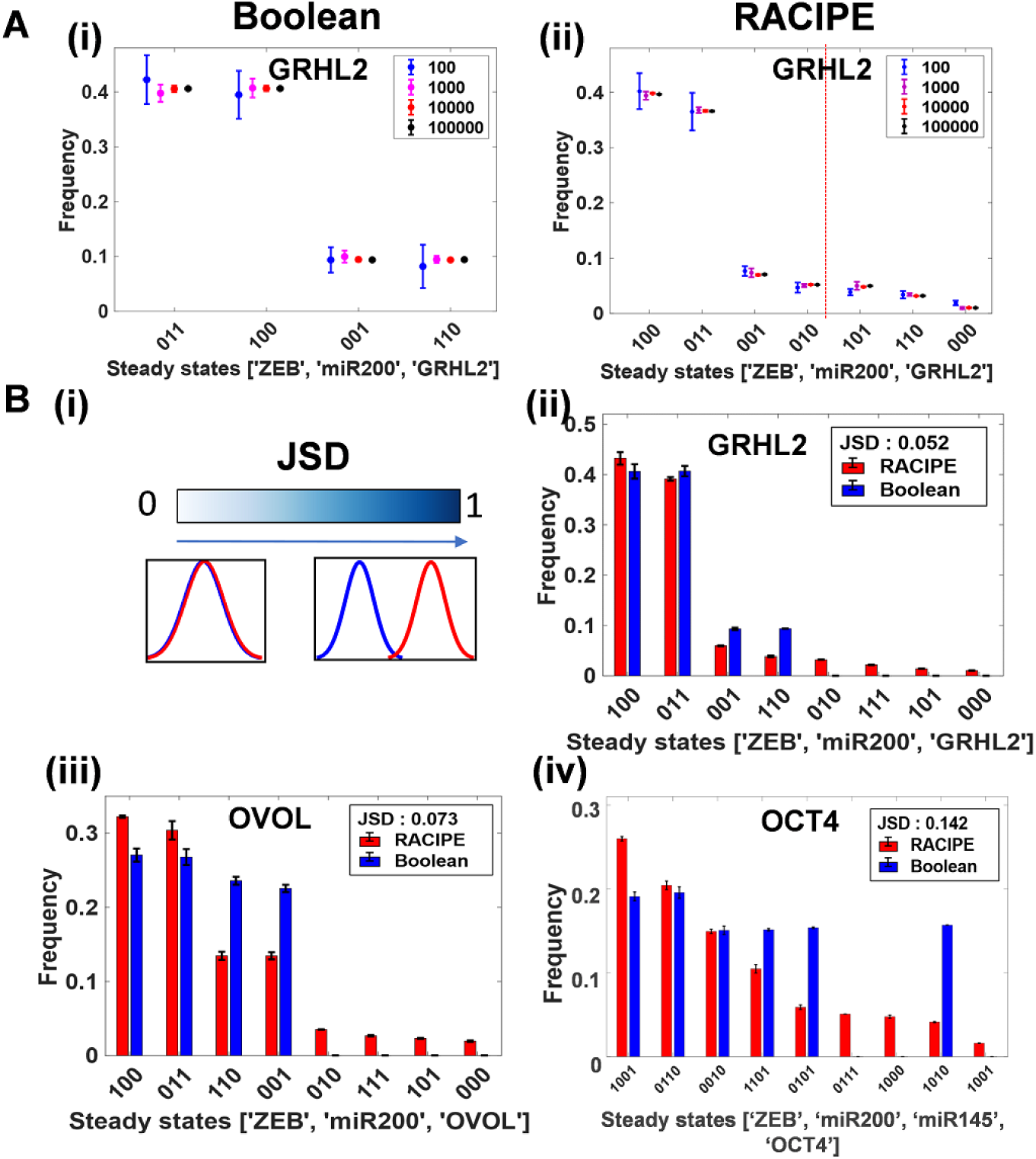
Comparing the outcomes of discrete (Boolean) and continuous (RACIPE) modelling frameworks. A. Quantitative convergence of the state frequency landscape for different number of initial conditions in Boolean analysis (left) and different number of parameter sets randomly sampled from the parameter space for RACIPE (right) respectively. Error bars represent the mean SD of the corresponding frequencies obtained by n = 3 independent simulations. B. (i) Demonstration of JSD between two given probability distributions; JSD ranges from 0 to 1. (ii)-(iv) Comparison of phenotypic frequency distributions for different EMP networks, as obtained from Boolean and RACIPE.

GRHL2 and OVOL1/2 are reported to play similar roles in inducing MET and/or inhibiting EMT [27, 31, 32]. Thus, epithelial (ZEB=0, miR-200=1, OVOL=1) and mesenchymal (ZEB=1, miR-200=0, OVOL=0) phenotypes are seen consistently as the most predominant ones in the RACIPE and Boolean model results for the SNAIL/miR-200/ZEB/OVOL network as well (**Fig S1A,i**). Since GRHL2 can self-activate [33] and OVOL can self-inhibit [34], we studied networks with GRHL2 self activation (referred to as GRHL2wa) and with and without OVOL self-inhibition (OVOL and OVOLsi respectively) (**Fig 1B**). In both Boolean and RACIPE, we observed that epithelial and mesenchymal states were the highest frequency phenotypes across these 3 cases (**Fig S1A,iii-iv**).

For the SNAIL/miR-200/ZEB/OCT4/miR-145 network (hereafter referred to as ‘OCT4’ network; 5 nodes, 10 edges), a maximum of 2^4^ = 16 stable steady states are possible (value of each node = 0 or 1; SNAIL is an input to the circuit). Boolean analysis across different numbers of initial conditions identified six out of the 16 possible states as stable steady states. The most predominant phenotypes were (ZEB=0, miR-200=1, miR-145=0, OCT4=0) and (ZEB=1, miR-200=0, miR-145=1, OCT4=1) which can be mapped on to epithelial and mesenchymal states correspondingly. Results obtained via RACIPE analysis are qualitatively consistent with those from Boolean model; with some additional but less frequent states identified via RACIPE (**Fig S1A,ii**). Similar consistency is observed when comparing the phenotypic distributions for SNAIL/miR-200/ZEB/miR-34/NRF2/KEAP1 network (hereafter called as ‘NRF2 network’; **Fig S1C**)

Next, for each of these different EMP networks, we quantified the difference between the phenotypic distributions obtained via RACIPE and Boolean models, using an information theory metric known as the Jensen-Shannon Divergence (JSD). JSD measures the dissimilarity between two given probability distributions and varies between 0 and 1 [35]; the larger the JSD, the more dissimilar or further apart are the two frequency distributions (**Fig 2B,i**). JSD for Boolean vs. RACIPE solutions for the EMP networks modelled here varies between 0.05 to 0.27 (**Fig 2B; S1B; S1C**), suggesting a good quantitative agreement between the two methods. Thus, these results indicate that the phenotypic distributions enabled by these EMP networks are largely a feature of the underlying network topology rather than of specific kinetic parameters.

### 2.2. Quantifying the effect of edge perturbations on phenotypic distributions of EMP networks via JSD

To characterize the effects of network topology on phenotypic distributions further, we made changes to the topology in the form of single-edge perturbations and quantified the impact of these edge perturbations on the phenotypic distributions obtained from various EMP networks. An edge perturbation can be one of the following: a) deleting an edge, b) adding a (hypothetical) edge, and c) changing the sign of an edge (i.e. from activation to inhibition or *vice-versa*). For a network with N nodes and E edges, there can be E edge deletions, (^*N*^*C*_2_ − *E*) additions, and E changes in edge sign. Thus, for the ‘wild-type’ (WT) SNAIL/miR-200/ZEB/GRHL2 network, 31 such perturbations are possible (**Table S1**), each of which will generate a new network topology. For every perturbation, we simulated the new network using both RACIPE and Boolean models and obtained the two corresponding phenotypic distributions. For the 32 distributions (31 perturbed + 1 ‘wild-type’) obtained via Boolean models, we then calculated the JSD between every two phenotypic distributions to identify perturbations that can drastically alter the phenotypic landscape. The network where the link from ZEB to GRHL2 was changed from inhibition to activation/excitation (*ZEB-GRHL2_2-1*) had the highest JSD from all remaining 31 networks (**Fig 3A**). RACIPE models, in addition to *ZEB-GRHL2_2-1* identified another perturbation which stood out relative to others – the deletion of the inhibitory link from ZEB to miR-200 (*Zeb-miR200_2-0*) (**Fig 3B**). Similar analysis for the NRF2 network using Boolean analysis identified two key perturbations while RACIPE analysis identified two additional ones (**Fig S2A**).

**Figure 3:**
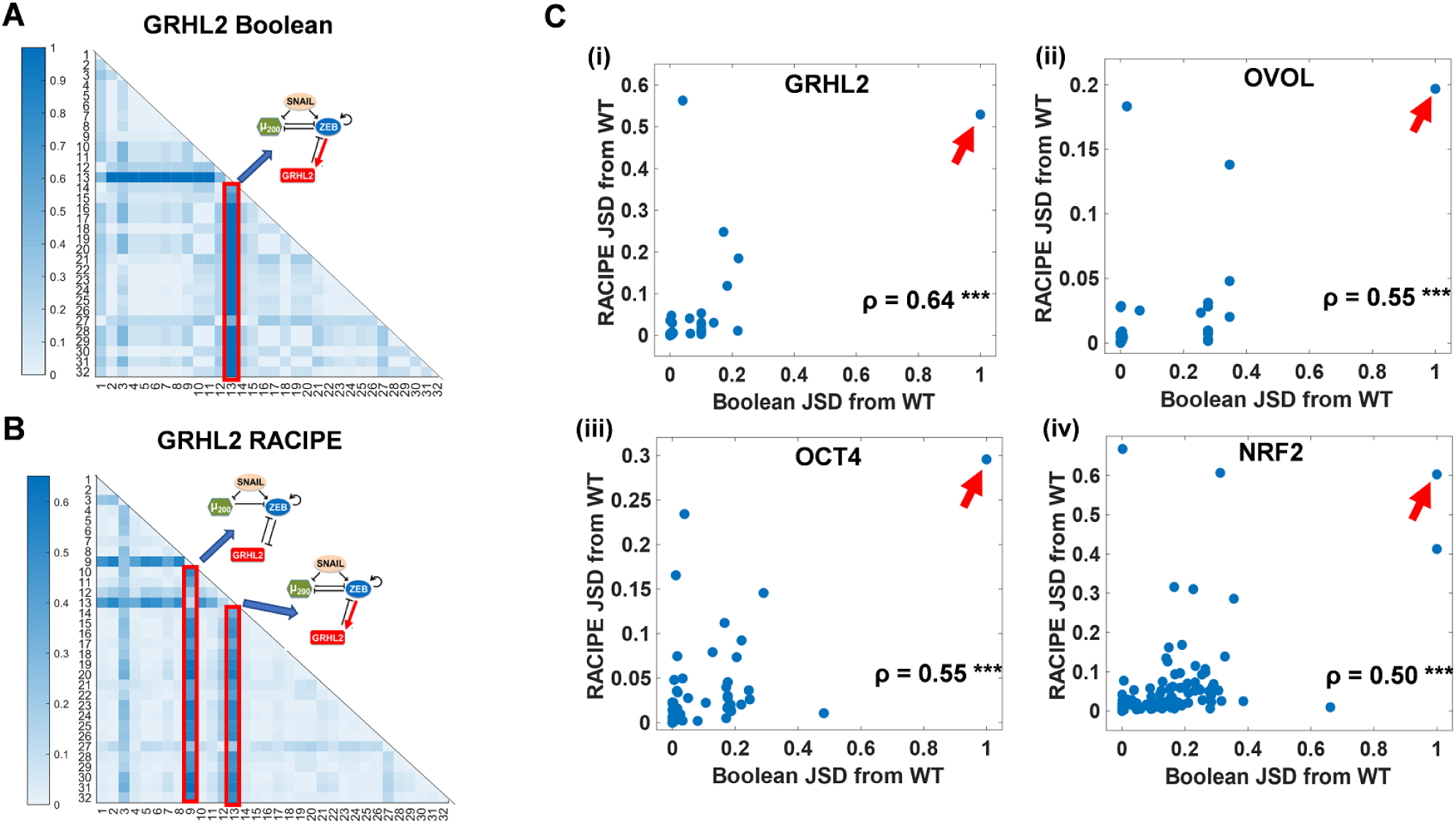
Quantifying the effect of single-edge perturbations on network behaviour landscapes. Heatmap of the JSD between steady state distributions of all perturbations for the GRHL2 network from each other, obtained from A. Boolean and B. RACIPE. Color bar shows the value of JSD. Each number (1-31) represents a particular perturbed network except for number 2, which represents ‘wild-type’ (**Table S1**). The perturbation highlighted in red has highest JSD from WT and most other perturbations. C. Scatter plots of JSD between the steady state distributions of a perturbed network from WT as obtained via RACIPE vs. as obtained via Boolean. Each dot in a plot represents a perturbed topology for the EMP networks – (i) GRHL2, (ii) OVOL, (iii) OCT4 and (iv) NRF2. The strongest perturbation identified by both Boolean and RACIPE is highlighted by the arrow. Spearman correlation coefficients (*ρ*) are reported; ***: *p* < 0.001

Further, we compared JSD between perturbed and ‘wild-type’ networks, calculated via RACIPE and Boolean analysis (**Fig 3C, S2B**). While the values of JSD were different for Boolean and RACIPE methods, a positive correlation was observed between the JSD values across all six EMP networks considered here. Moreover, the strongest perturbation obtained via both methods showed concordance (highlighed by the arrows in **Fig 3C and S2B**). These results further emphasize the role of network topology in phenotypic distributions generated by EMP networks.

### 2.3. Correlation between JSD and phenotypic plasticity is not consistently significant across EMP networks

Next, we investigated whether the perturbed networks which are farthest from the ‘wild-type’ network (i.e., having the highest JSD) are the ones with reduced phenotypic plasticity as well. Phenotypic plasticity is the ability of cells to sample multiple phenotypes and to switch from one phenotype to another, spontaneously or under external factors. The definition of phenotype in the present context can either be a stable steady state identified by mathematical simulations of EMP networks or a biological cell-state obtained by classifying the steady states based on marker expression levels. Hence, we define phenotypic plasticity in two different ways using RACIPE output. For every randomly chosen parameter set, RACIPE simulates the system with multiple, randomly chosen, initial conditions. For some parameter sets, all chosen initial conditions converge to one stable state, while in others, multiple steady states (multistability) may be allowed. Thus, plasticity score 1 (PS1) is defined as the fraction of parameter sets that enable multistability (**Fig 4A**). The definition of plasticity score 2 (PS2) is more biology-centric. We first define the ‘phenotype’ of a given steady state based on the discretized expression levels of canonical epithelial and mesenchymal markers – miR-200 and ZEB respectively [29, 36, 37]. This allows for the identification of various phases (combinations of co-existing steady states), such as the co-existence of epithelial and mesenchymal states {E,M} for instance. PS2 is the fraction of parameter sets which allow multiple phenotypic states, i.e. multi-stable phases (**Fig 4A**). For a given network, we calculated PS1 and PS2 scores for the ‘wild-type’ and perturbed topologies; a comparison of these two metrics revealed a strong positive correlation across all 6 networks (**Fig 4B, S3A**). The perturbed networks had both increasing as well as decreasing effect on phenotypic plasticity (**Fig 4B, S3A**).

**Figure 4:**
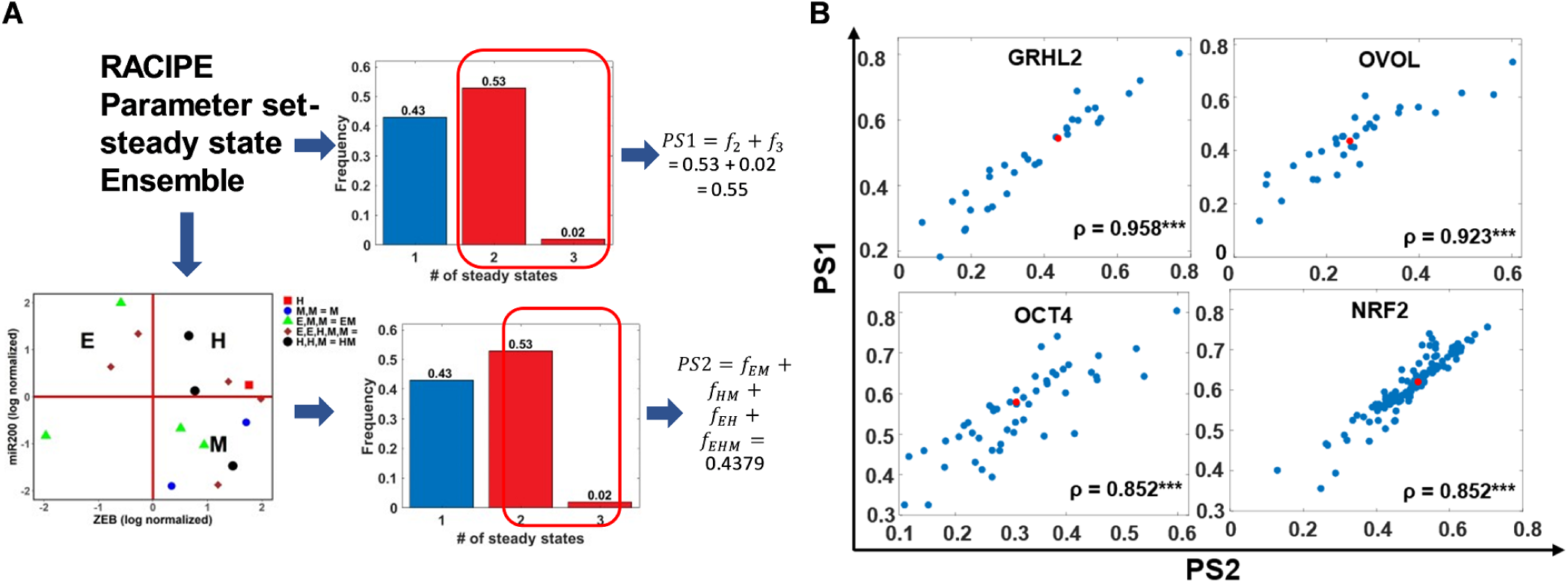
Metrics for quantifying phenotypic plasticity for the case of single-edge perturbations. A. Two definitions or plasticity: PS1 (top-right) is defined as the fraction of multi-stable parameter sets identified by RACIPE. PS2 (bottom) is calculated after ZEB and miR200 expression levels are used to classify steady states obtained from each parameter set (denoted by different colors) into Epithelial (E) – (high miR-200, low ZEB) / Hybrid (H) – (high miR-200, high ZEB) / Mesenchymal (M) – (low miR-200, high ZEB) phenotypes. Parameter sets are then characterized as monostable or multi-stable based on the phenotypic states they sample. PS2 is defined as the fraction of parameter sets giving rise to multiple phenotypes. B. Scatter plot of PS1 vs. PS2 for different EMP networks – WT (colored red) and perturbed (colored blue; single-edge perturbed: edge deletion, edge nature change and edge additions) ones. Spearman correlation coefficients (*ρ*) are denoted; ***: p <0.001

Further, we checked whether the topologies with the highest JSD from the ‘wild-type’ network led to a decrease or an increase in PS2 scores. We did not observe any significant overlap of the network topologies with the highest JSD vs. those with the highest or the lowest PS2 scores. This lack of trend was seen across all six networks considered (**Fig 5A**). Further, a scatter plot between JSD from the ‘wild-type’ network and fold-change in PS2 scores relative to the ‘wild-type’ was plotted. While some networks showed a negative correlation, others had no significant correlation between these two metrics (**Fig 5B; S3B**) Similar results were obtained for analysis done using PS1 scores for these perturbed networks (**Fig S3C**). Together, these observations suggest that JSD is not a good predictor of change in phenotypic plasticity.

**Figure 5:**
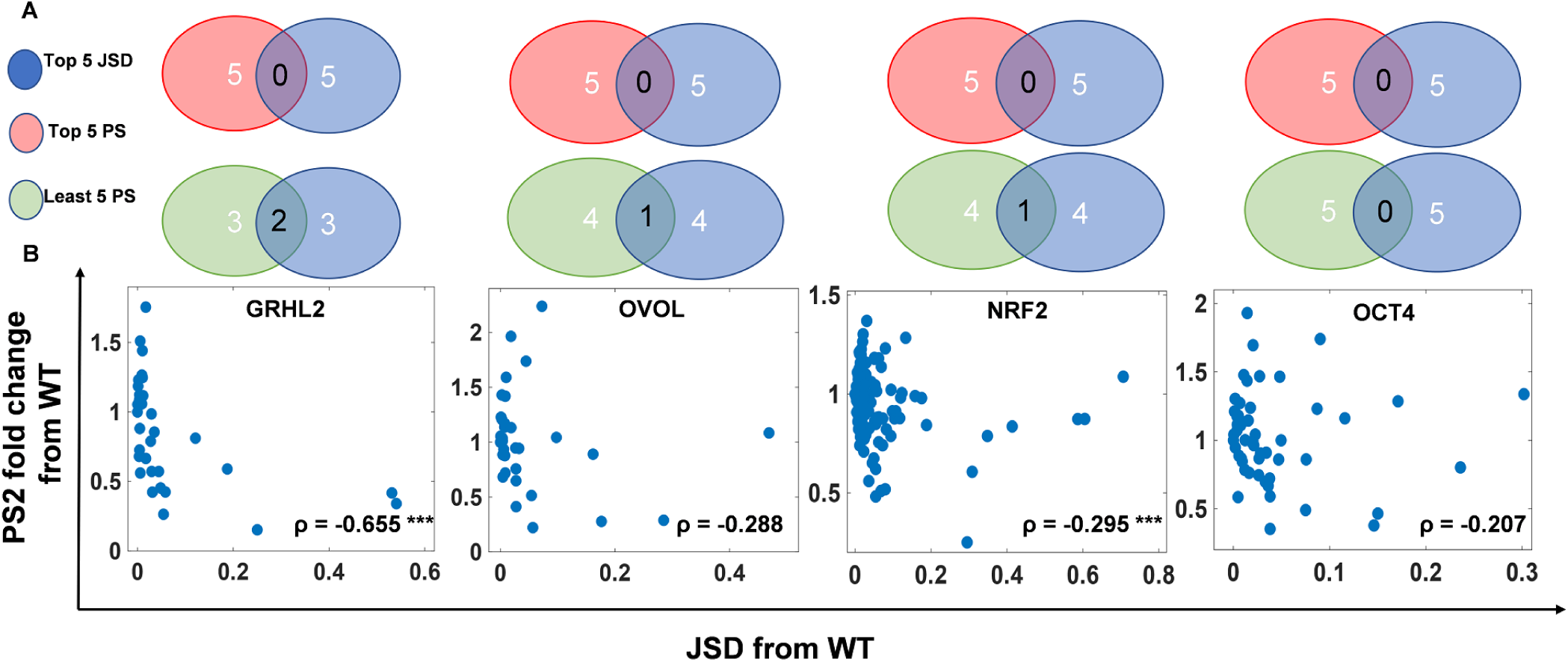
Correlation between JSD and phenotypic plasticity across EMP networks. A. Venn diagrams showing the extent of overlap among the perturbed networks that have the highest JSD from the ‘wild type’ network and those with the highest or lowest fold change in plasticity scores (PS2). EMP networks shown are (from left to right): GRHL2, OVOL, NRF2 and OCT4. B. Scatter plots for respective EMP networks. Each blue dot in a scatter plot is a perturbed network topology corresponding to that EMP network. Spearman correlation coefficient (*ρ*) are denoted; ***: p<0.001

### 2.4. Number of positive feedback loops in EMP networks correlates positively with phenotypic plasticity

Next, we revisited the network topologies with the highest JSD from the ‘wild-type’ network to identify any topological signatures and observed that all of them were disrupting an overall positive feedback loop in the ‘wild-type’. Here, the ‘overall’ sign of a loop is defined by the product of signs of edges (positive for activation, negative for inhibition) that form a cycle/loop; thus, a mutually inhibitory loop between two molecular players is effectively a positive feedback loop. In the GRHL2 network, the deletion of ZEB to miR-200 inhibitory link (*ZEB-miR200_2-0*) disrupted the mutually inhibitory loop between ZEB and miR-200. Similarly, converting the inhibitory link from ZEB to GRHL2 to being excitatory (*ZEB-GRHL2_2-1*) disrupted the mutually inhibitory feedback loop between ZEB and GRHL2 (**Fig 3A**). In the NRF2 network, converting the inhibitory link from ZEB to E-cadherin to being excitatory (*ZEB-Ecad_2-1*) disrupted the mutually inhibitory loop between ZEB and E-cadherin, and converting the inhibitory link from miR-200 to NRF2 to excitatory (*miR200-NRF2_2-1*) disrupted the overall positive feedback loop formed by miR-200, KEAP1, NRF2 and SNAIL (**Fig S2A**).

Previous analysis for simpler two-node networks has shown that mutually inhibitory and mutually excitatory loops (hence, both being overall positive loops) can lead to multistability which may drive phenotypic plasticity [38, 39]. Such networks are typically observed underlying the cell-fate decisions during embryonic development [40]. Similar observations have been made for miR-200/ZEB feedback loop in driving trans-differentiation through EMP [36, 41]. Therefore, we further inquired whether phenotypic plasticity can be correlated with the total number of positive feedback loops in a given network. We counted the number of positive feedback loops/cycles for all ‘wild-type’ and perturbed topologies for all six EMP networks (see Materials and Methods for description). First, taking GRHL2 network as a case study, we showed that decreasing the number of positive feedback loops by one (WT-1) reduced phenotypic plasticity (**Fig 6A**), while the reverse was true when the number of positive feedback loops was increased by one (WT + 1, **Fig 6A**).

**Figure 6:**
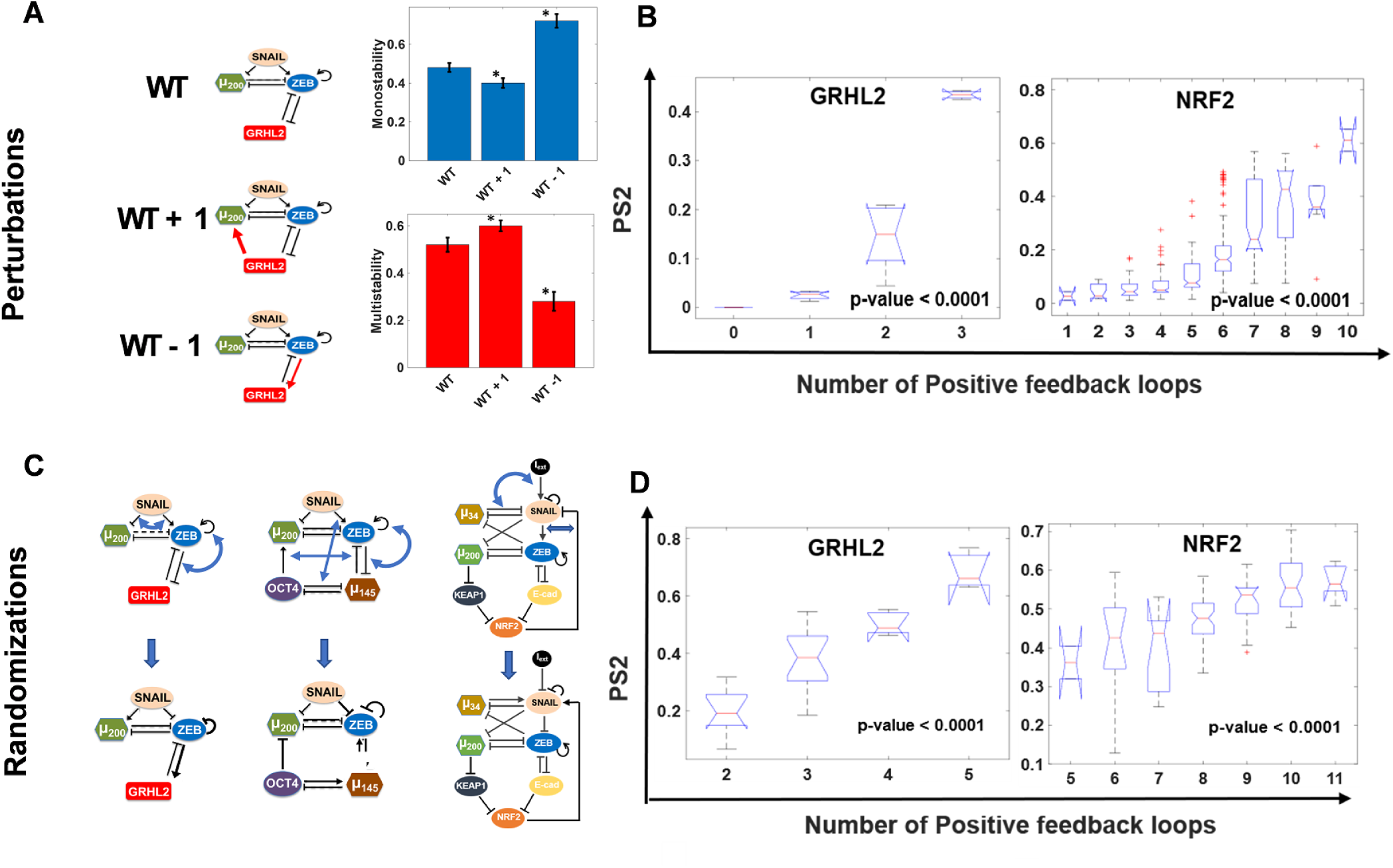
Effect of positive feedback loops on phenotypic plasticity in EMP networks. A. Demonstration of change in plasticity on altering the total number of positive feedback loops in GRHL2 network. Frequency of multi-stable parameters (PS2) increases as compared to WT upon the addition of positive feedback loop (GRHL2-miR200-ZEB-GRHL2) to the network (WT + 1) and reduces upon reduction of positive feedback loops by changing the GRHL2-ZEB-GRHL2 cycle to negative feedback (WT – 1). B. Boxplots of PS2 vs. number of positive feedback loops for all the perturbed networks for a given EMP network module. C. Demonstration of network randomization (the in-degree and out-degree of each node is preserved; however, the number of inhibitory/excitatory nodes arriving at or emerging from a node are not necessarily conserved). D. Same as B, but for randomized network topologies. p-value range for the one-way ANOVA test are mentioned on the plots.

Next, we compared the number of positive feedback loops in each perturbed network with the corresponding plasticity score, using one-way ANOVA. Indeed, the mean plasticity score is higher for the groups of networks with higher number of positive feedback loops; this trend is observed across all six EMP networks in a statistically significant manner for both plasticity metrics – PS1, PS2 (**Fig 6B; S4A-B**), suggesting a correlation between the number of positive feedback loops in an EMP network, and its ability to give rise to phenotypic plasticity. We also observed that the total number of feedback loops (i.e., positive feedback loops + negative feedback loops) in the networks themselves did not exhibit any significant and consistent effect on plasticity, further emphasizing the role of positive feedback loops specifically (**Fig S5**).

To test whether this observed correlation between plasticity scores and the number of positive feedback loops is specific to the network topology studied, we generated randomized topologies for each given network by swapping the edges in a given network (**Fig 6C**). This procedure preserves the in-degree and out-degree of each node in the network but can change the distribution of excitatory or inhibitory links arriving at (in-degree) or originating from (out-degree) a given node. Thus, for a given EMP network, such randomization can generate various network topologies with varying number of net positive feedback loops. For each randomized network topology, we calculated PS1 and PS2. Reinforcing our results to perturbed networks, we observed a positive trend between plasticity and positive feedback loops (**Fig 6D; S4C-D**), strengthening our hypothesis that the number of positive feedback loops in a given EMP network is a good predictor of phenotypic plasticity.

### 2.5. JSD, Positive feedback loops, phenotypic plasticity and scalability of the corresponding trends

While the plasticity scores correlated positively with the number of positive feedback loops, there was heterogeneity in plasticity scores for a set of network topologies having the same number of positive feedback loops (**Fig 6B,D; S4**), indicating the presence of other factors in addition to positive feedback loops that determine phenotypic plasticity. Because a weak and inconsistent negative correlation was observed between JSD and plasticity scores (**Fig 5; S3B-C**), we asked if JSD can contribute to explaining heterogeneity in plasticity.

For each network topology obtained by perturbing or randomizing the corresponding EMP network, we projected the plasticity score on a 2-D plot of its JSD from the wild-type and the number of positive feedback loops (**Fig 7A,i; 7B,i; S6**). These plots convey that JSD did not resolve the heterogeneity in plasticity scores of network topologies with the same number of positive feedback loops. To quantify this observation, we discretized JSD into ranges of values and calculated the correlation between positive feedback loops and plasticity for a range of JSD and correlation between JSD and plasticity for a fixed number of positive feedback loops.

**Figure 7:**
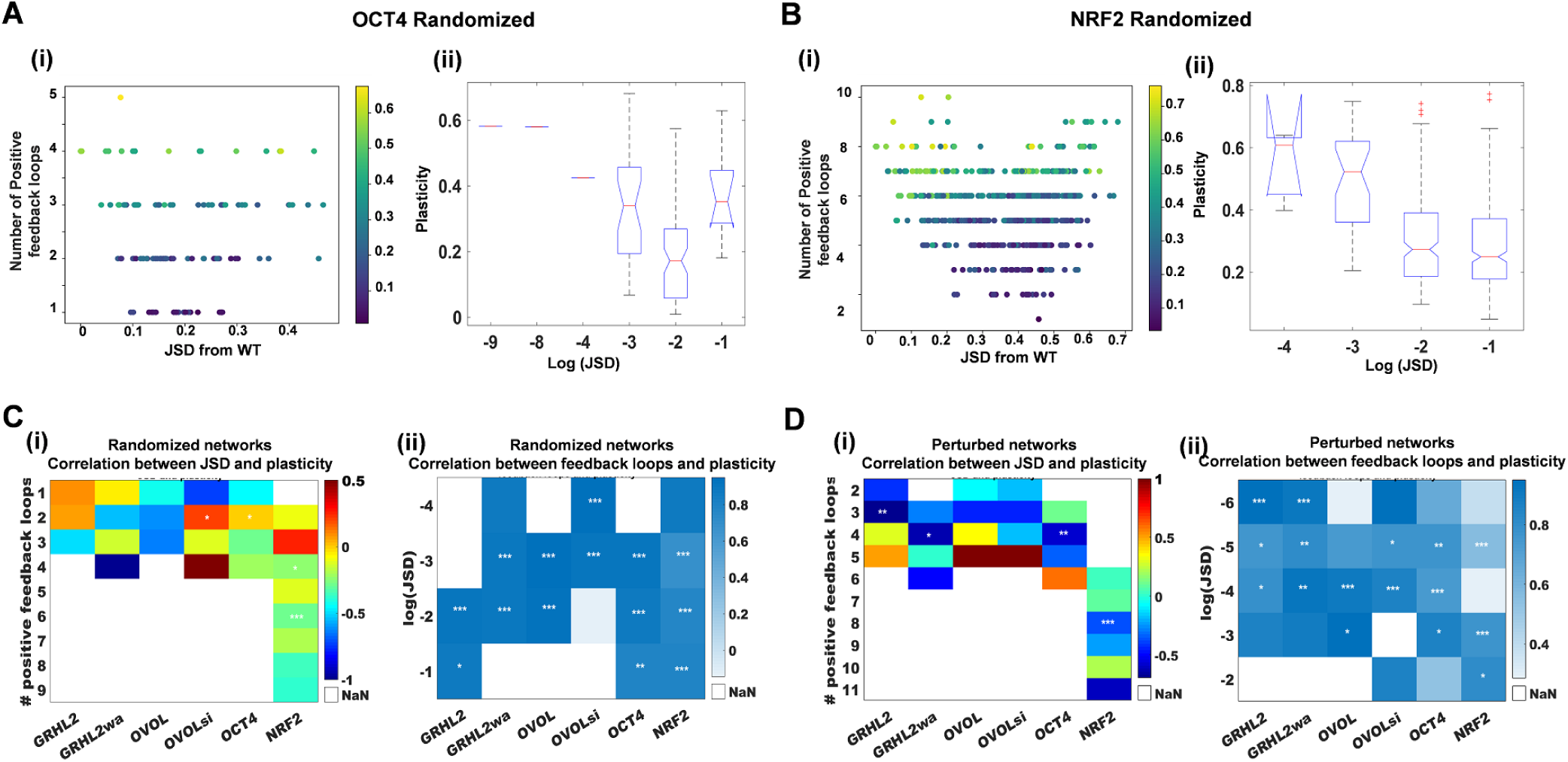
Correlation between JSD, plasticity, and number of positive feedback loops. A. (i) PS2 for every randomized network for OCT4 module plotted as a function of its JSD from the WT network and number of positive feedback loops. Colorbar shows PS2 scores. (ii) Boxplot for PS2 scores vs. range of JSD values for all networks plotted in (i); p-value corresponds to one-way ANOVA test. B. Same as A but for NRF2 module C. (i) For each EMP module, all randomized networks are sub-categorized based on number of positive feedback loops. Each cell denotes correlation coefficient between JSD from WT and plasticity score fold change, wherever applicable (i.e. number of samples corresponding to that network and feedback loop count > 3. The white boxes (NaN) represent the cases with number of samples less than 3). Spearman correlation coefficient shown using color bar; significance represented as: *: *p* < 0.05, **: *p* < 0.01, * * *: *p* < 0.001 (ii) For each EMP module, all randomized networks are classified based on range of JSD values. Each cell denotes correlation coefficient between plasticity score fold change and number of positive feedback loops. D. Same as C but for perturbed network for each EMP module.

The categorized JSD values maintained the lack of consistent correlation with plasticity (**Fig 7A,ii; 7B,ii**) as seen earlier (**Fig 5; S3B-C**). The lack of correlation was not ameliorated by grouping of the network topologies by the number of positive feedback loops for either perturbed or randomized topologies. The correlation coefficient varied from −1 to 1 even within a given EMP network with many instances being statistically insignificant (**Fig 7C,i; 7D,i**). On the other hand, when these network topologies were segregated based on JSD, plasticity scores and number of positive feedback loops were positively correlated across most ranges of the JSD values and across the six EMP networks (**Fig 7C,ii; 7D,ii**). Together, these results strongly support positive feedback loops as good predictor of phenotypic plasticity and suggest that phenotypic frequencies may not be useful in measuring phenotypic plasticity.

To test the scalability of these results, we analysed two larger EMP networks: EMT RACIPE [25] (22 nodes, 72 edges, **Fig 8A,i**) and EMT RACIPE2 [24] (26 nodes, 101 edges **Fig 8A,ii**), and all single-edge deletions and edge nature change (activation to inhibition and vice-versa) perturbations (n=144 and 202 respectively) for these networks. Given the network complexity, it becomes increasingly challenging to unquely associate mathematically observed stable steady states with biological phenotypes. Hence, we used the generic definition of phenotypic plasticity (PS1). Similar to the smaller networks, the correlation observed between change in the number of positive feedback loops due to single-edge perturbations and the corresponding fold change in plasticity from WT was positive and significant (**Fig 8B,ii; 8C,ii**). Similarly, the correlation between JSD and plasticity was weak and insignificant (**Fig 8B,i; 8C,i**). Furthermore, the 2-D plots projecting plasticity scores onto JSD (x-axis) and change in positive feedback loops (y-axis) showed no distinct patterns (**Fig 8B,iii; 8C,iii**), supporting the conclusion that JSD cannot reliably predict changes in phenotypic plasticity.

**Figure 8:**
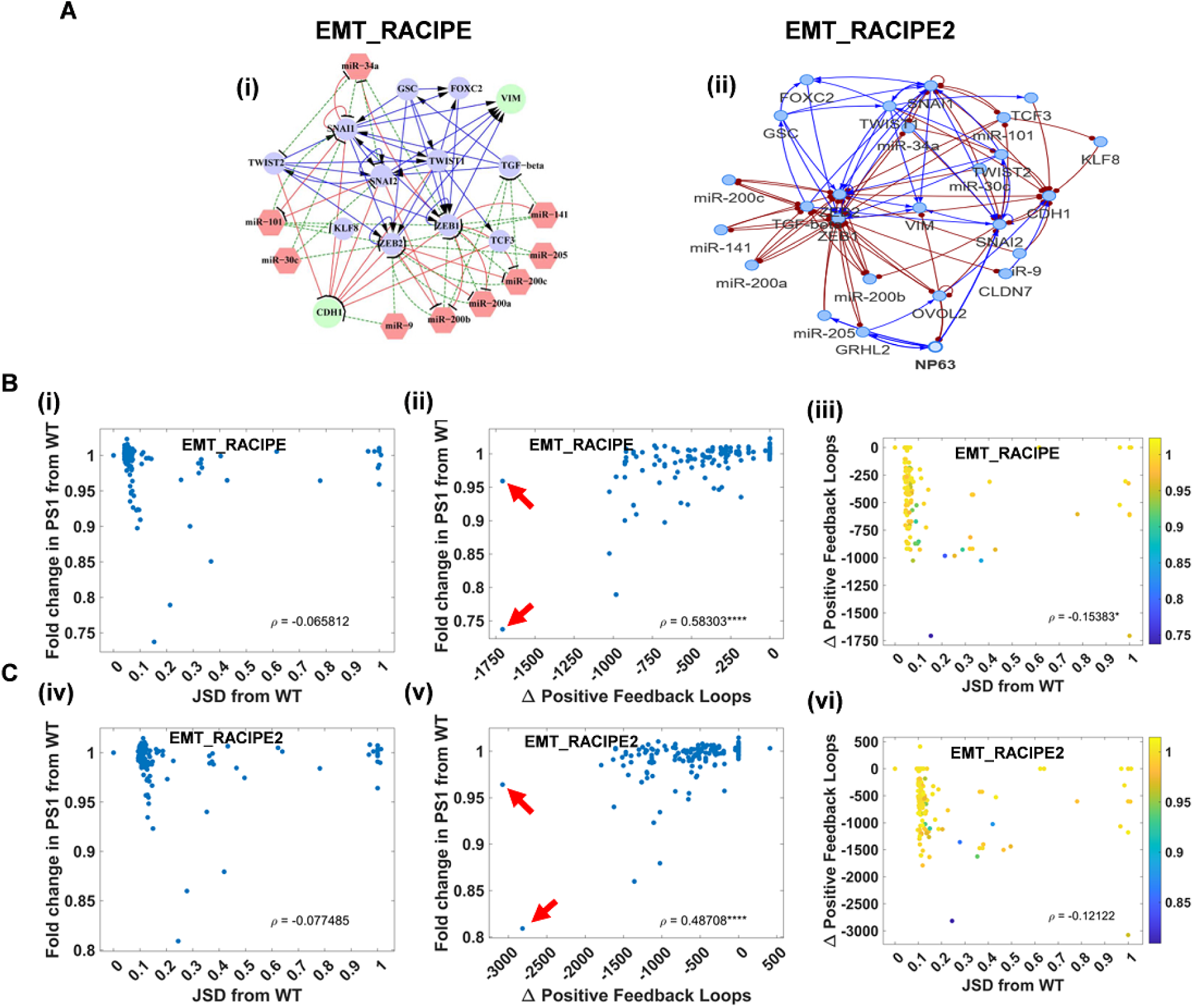
Analysis of 22 and 26 node networks. A. (i) The 22 node EMP network (EMT RACIPE). (ii) The 26 node EMP network (EMT RACIPE2) B. EMT RACIPE (22 node, 72 edges). (i) Scatter plot between PS1 and change in total number of positive feedback loops of all the perturbations. (ii) Scatter plot between PS1 and JSD of all perturbations from WT. (iii) Scatter plot of JSD from WT and change in positive feedback loops for each perturbed network. Color code denotes absolute value of PS1 for a given perturbed network topology. C. Same as B but for EMT RACIPE2 (26 node, 101 edges).

Owing to the huge number of positive feedback loops in these two networks, we were able to observe a saturation in the effect of the number of positive feedback loops on plasticity. Interestingly in these larger circuits, single-edge perturbations are capable of disrupting hundreds of positive feedback loops. The maximum number of positive loops disrupted by a single edge is approximately 1750 in EMT RACIPE and 3000 in EMT RACIPE2, which account for more than 50% positive feedback loops in wild-type topology. Furthermore, two perturbations were observed in both networks that reduce the positive feedback loops by half or more (pointed by red arrows), but only one of them significantly reduces plasticity. These observations, together with the observation that in smaller networks, multiple networks with the same number of positive feedback loops can have different levels of plasticity **(Fig 6; Fig S4)** point towards additional network topological features that may contribute to determine the phenotypic plasticity resulting from EMP networks.

## 3. Discussion

Our ability to target phenotypic plasticity is limited by the understanding of its dynamics in tumors and the identification of tumor-specific molecular mechanisms driving it. While various transcriptional and epigenetic networks have been uncovered underlying phenotypic plasticity; how do these networks give rise to the co-existence of various drug-tolerant states, and the contribution of these different states to the minimal residual disease remains elusive [42]. Recent surge in our understanding of the dynamics of EMP has elucidated how the underlying EMP regulatory networks can give rise to various phenotypes/cell states along the spectrum of EMP, and how these different sub-populations may co-exist in a tumor and collaborate to drive tumor aggressiveness [43]. Nonetheless, targeting EMP in the clinic to observe a major reduction in metastasis still remains a challenge, because of lack of understanding of the precise spatio-temporal regulation of EMP involved during metastasis. Inhibiting one arm of EMP – say EMT – might actually promote MET and help colonization; and inhibiting MET may facilitate more dissemination [7]. Moreover, inhibiting only EMT or MET may drive the cells into one or more hybrid E/M phenotypes that are considered to be the ‘fittest’ for metastasis [13]. Therefore, while targeting EMP is important for restricting metastasis and therapy resistance, how to achieve that remains an unsolved challenge.

Our results present a computational platform to identify specific inhibitors for EMP using a network-level approach. We have simulated various networks identified to underlie EMP using different modeling strategies, to dissect the design principles underlying those networks; and suggest how perturbing those networks may prevent the ability of cells to switch back and forth among the E, M and hybrid E/M phenotypes. Our analysis predicts that reducing the total number of positive feedback loops in the EMP network can restrict plasticity. A recent experimental study offers preliminary validation of this prediction, where disrupting the miR-200/ZEB mutually inhibitory (hence, an overall positive feedback) loop via CRISPR led to significantly reduced metastasis in vivo [41]. This feedback loop has been identified as a key regulator of EMP through extensive experimental and computational analysis [29, 36, 37]. Mathematical modeling for this loop has predicted how clonal cells responding to the same EMT-inducing signal can display different phenotypes due to the emergent multistability (the co-existence of multiple steady states/ phenotypes), a prediction which was validated experimentally via single-cell analysis of EMP [41]. Various other EMP networks that have been mathematically studied have included various other direct or indirect positive feedback loops such as ZEB1/GRHL2 [30], ZEB1/ ESRP1 [44], ZEB1/miR-1199 [45], or miR-34/SNAIL [46].

Different modeling frameworks have been used to investigate the dynamics of EMP, depending on the size of network. While small-sized networks have typically been modelled via continuous approaches [47, 48, 9, 36, 49, 50], larger networks have been modelled via discrete Boolean approaches due to lack of available kinetic parameters [24, 45, 51, 52]. While continuous models provide a more quantitative mapping of system dynamics but require many kinetic parameters that can become experimentally intractable, Boolean modeling approaches provide a good estimate of qualitative behavior of a biochemical system without requiring a large set of parameters [53], but are limited in terms of characterizing dynamic properties such as phenotypic plasticity and state transition rates. Thus, various efforts have been made to compare the dynamics of Boolean vs. continuous models and to integrate their strengths, particularly for capturing steady state distributions for smaller biological networks [54, 55, 56].

Here, we compare the phenotypic distributions obtained for various EMP networks using Boolean formalism [24] and using RACIPE [25]. While Boolean tries to capture the phenotypes that can non-parametrically be observed in a network; RACIPE, by simulating networks for a large number of parameter sets, tries to capture the effect of parameteric variability observed in cell populations from one or multiple individuals on the phenotypes. Both methods infer network behavior as a function of the topological information, unbiased by any specific parameter set. Hence, the qualitative and semi-quantitative agreement seen for Boolean and RACIPE models, across six EMP networks, enable us to understand the dynamics of EMP driven by network topology instead of specific kinetic parameter sets in a given cell/population. Furthermore, we could identify perturbations to the network topologies that affetced the phenotypic distributions significantly across parameter sets. Thus, our method to identify network-topology based predictions to inhibit EMP may provide an avenue to overcome a major bottleneck in targeted therapy — inter-individual variability in response. Moreover, through generating a larger number of randomized networks where the in-degree and out-degree of each node in the network was preserved, we showed that the phenotypic distributions and plasticity scores (PS1, PS2) obtained are specific to the particular topology of the networks regulating EMP. These results suggest evolutionary design principles of EMP networks that may have been optimized to induce EMP as/when needed during development/tissue regeneration, and stably maintain homeostatic differentiated phenotypes.

Intriguingly, the change in phenotypic plasticity, defined by both the plasticity scores does not correlate with JSD of phenotypic distributions. One possible reason underlying this perceivably confounding result can be that the JSD only computes the distance between two steady state distributions; it does not capture information about whether the phenotypes can switch among themselves. (Spontaneous) phenotypic switching is facilitated by multi-stable phases, i.e. the co-existence of more than one stable steady state, for a given parameter set. Our results that the number of positive feedback loops in a given network determines the extent of phenotypic plasticity is reminiscent of reported connection between positive feedback loops and plasticity in other aspects of cancer too, where mutually inhibitory loops between two ‘master regulators’ drive phenotypic switching [57, 58, 59, 60]. Future efforts should aim at identifying which links(s) in the network to disrupt to cause maximal change in plasticity, because not every positive feedback loop may be equally likely to lead to multi-stability [61, 62]. Moreover, for networks with the same number of feedback loops, the plasticity scores varied over a range, thus, identifying other predictors of plasticity based on network topology will be valuable. As an attempt to identify such complementary predictors, we investigated if JSD coupled with number of feedback loops can lead to isolate networks with highest plasticity, but no clear improvement in the trend was observed, eliminating JSD as a predictor of plasticity either individually or in combination with feedback loops.

Our results for the 22 and 26-node EMP network via RACIPE identified edge deletions that can reduce the positive feedback loops by half and have a significant impact on the plasticity. As more comprehensive networks representing the underlying biology are studied, RACIPE becomes too computationally expensive, and hence network theory-based measures to identify the feedback loop that, when disrupted, can have maximal effect in curbing plasticity would be valuable for future therapeutic applications.

It should be noted that while the existence of greater than or equal to two stable states is essential for phenotypic switching, we need to take into account the relative stability of these states, which determines the cellular transition rates from one state to another. In the context of EMP, hybrid epithelial mesenchymal states are known to be relatively less stable as compared to pure epithelial or pure mesenchymal states and hence are highly plastic [63]. Hybrid phenotypes are also associated with increased metastatic potential [17, 64]. Thus, future efforts to restrict EMP bidirectionally should consider these state-specific traits to identify and rank various possible interventions to the EMP network topology.

Most of the targeted therapies in oncology target on disrupting a node in the network. Inevitably, most cells can identify ‘escape’ routes by navigating various dimensions of the phenotypic plasticity landscape. Our results present an alternative and unorthodox mechanism to restrict the emergence of metastasis and drug resistance – breaking the feedback loops, i.e. targeting a link instead of a node, involved in phenotypic plasticity. Disrupting these feedback loops – the cornerstone of phenotypic plasticity – can restrict the ability of cancer cells to adapt to various therapeutic attacks and limit tumor aggressiveness.

## 4. Methods

### 4.1. RAndom CIrcuit PErturbaiton (RACIPE)

RACIPE [25, 65] is a tool that simulates transcriptional regulatory networks (TRNs) in a continuous manner. Given a TRN, it constructs a system of Ordinary Differential Equations representing the network. For a given node *T* and a set of input nodes *P*_*i*_ and *N*_*j*_ that activate and inhibit *T* respectively, the corresponding differential equation takes the following form:

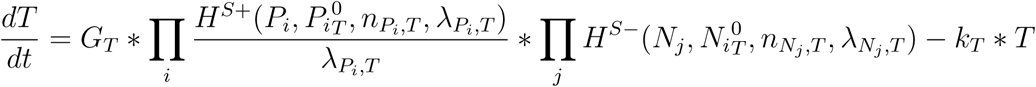

Here, *T, P*_*i*_ and *N*_*j*_ represent the concentrations of the species. *G*_*T*_ and *k*_*T*_ denote the production and degradation rates, respectively. 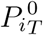 is the threshold value of *P*_*i*_ concentration at which the non-linearity in the dynamics of *T* due to *P*_*i*_ is seen. *n* is termed as hill-coefficient and represents the extent of non-linearity in the regulation. *λ* represents the fold change in the target node concentration upon over-expression of regulating node. Finally, the functions *H*^*S*+^ and *H*^*S*−^ are known as shifted hill functions [36] and represent the regulation of the target node by the regulatory node. The hill shift function takes the following form:

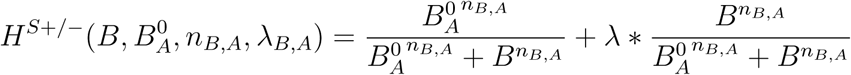

Note that, for high values of the regulatory node concentration, *H*^*S*+/−^ approaches λ.

For the model generated in this way, RACIPE randomly samples parameter sets from a predefined set of parameter ranges estimated from BioNumbers [66]. The ranges as reported by Huang et al [65] are as follows:

**Table 1:** Parameter ranges used by RACIPE.

| Parameters | Minimum | Maximum |
| --- | --- | --- |
| Production Rate ($G$) | 1 | 100 |
| Degradation Rate ($k$) | 0.1 | 1 |
| Fold Change (Inhibition $\lambda$) | 0.01 | 1 |
| Fold Change (Activation $\lambda$) | 1 | 100 |
| Hill Coefficient $n$ | 1 | 6 |
| Threshold | The ranges depend on inward regulations - half functional rule |  |

At each parameter set, RACIPE integrates the model from multiple initial conditions and obtains steady states in the initial condition space. The output, hence, comprises of the collection of parameter sets and corresponding steady states obtained from the model. For the current analysis, we used a sample size of 10000 for parameter sets and 100 for initial conditions. The parameters were sampled via a uniform distribution and the ODE integration was carried out using Euler’s method of numerical integration.

### 4.2. Boolean simulations

For discrete analysis of our networks, the Boolean algorithm devised by Font-Clos et al., [24] was used. The nodes are updated asynchronously according to a majority rule such that the state of a node is set to 1 if the sum of activations to the node is more than the sum of inhibition and set to 0 if inhibition is more than activation. If inhibition and activations are equal, nodes are not updated. Steady state is said to have reached if the state of the network doesn’t change over time-steps. The input for this formalism is a set of 10000 initial conditions, which are randomly sampled from all possible states of the system and corresponding steady states.

### 4.3. Discretization of RACIPE output and calculating the state frequency

For a given network with *i* = [1, *n*] nodes, the steady state expression levels of the nodes were normalized in the following way:

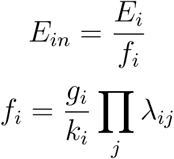

Where, for the *i*^*th*^ node, *E*_*in*_ is the normalized expression level, *E*_*i*_ is the steady state expression level, *f*_*i*_ is the normalization factor, *g*_*i*_ and *k*_*i*_ are production and degradation of the *i*^*th*^ node corresponding to the current steady state and *λ*_*ij*_ are the fold change in expression of *i* due to node *j* = [1, *n*]. The normalized expression levels of all steady states are then converted into z-scores by scaling about their combined mean:

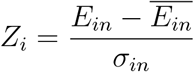

where 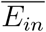 is the combined mean and *σ*_*in*_ is the combined variance.

The z-scores are then classified based on whether they are negative or positive into 0 (low) and 1 (high) expression levels respectively. Each steady state of the network is thus labelled with a string of 1’s and 0’s, discretizing the continuous steady state levels. We then calculate the total frequency of each discrete state by counting the occurrence in all the parameter sets. For parameter sets with n steady states, the count of each steady state is taken as 1/n, invoking the assumption that all the states are equally stable.

### 4.4. Quantitative convergence

To estimate the optimal sample size of parameter sets for RACIPE and that of initial conditions for Boolean models, all networks were simulated at different sample sizes in triplicates and the mean an variance of the steady state frequency distribution was calculated. 10000 was estimated as the ideal sample sizes for both methods as it was the smallest sample size for which the variance in steady state frequencies was minimum and the mean of the same was consistently similar to that of higher sample sizes.

### 4.5. Jensen-Shannon Divergence to measure distance between phenotypic distributions

To quantify the difference between two phenotypic distributions, an information theory metric, known as the Jensen-Shannon divergence (JSD) [35] was used, calculated for any two discrete frequency distributions *P* (*x*) and *Q*(*x*) as:

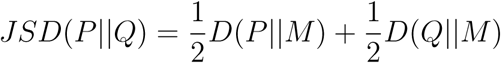

where 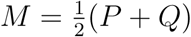 and *D* denotes the Kullback-Leibler divergence, defined as

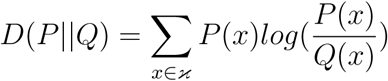

The metric, JSD, varies between 0 and 1 when the base 2 logarithm is used for calculation, with 0 indicating identical distributions and 1 indicating no overlap between the two distributions.The JSD calculations were done using the *JSD* function from *philentropy* package in *R 3.6*.

### 4.6. Calculating the number of positive feedback loops in a network

We estimated the number of feedback loops in the networks using the *networkx* module in *Python 3.7*, where a feedback loop is defined as a path traversed along the edges of a network that originates and ends at the same node. We then combine the nature of edges in each feedback loop to determine whether the given feedback loop is positive or negative. For example, in the OCT4 network (Fig 1B), ZEB-miR200-ZEB is a positive feedback loop, as it goes through 2 inhibitory edges. On the other hand, ZEB-miR145-OCT4-miR200-ZEB is a negative feedback loop, as the edges involved are inhibition-inhibition-activation-inhibition, in that order.

### 4.7. Statistical tests

All correlation analysis was done using Spearman correlation method using ‘cor’ function in MATLAB R2018b (Mathworks). The corresponding statistical significance values are represented by ‘*’s, to be translated as: *: p<0.05, **: p<0.01, ***: p<0.001. One-way ANOVA test was performed using anova1 function in MATLAB R2018b (Mathworks).

## 5. Author Contributions

MKJ and FFC designed the research; CLP and SZ analysed the data; KH, BS and BVS carried out the simulations; all authors discussed results and participated in the preparation of the manuscript.

## 7. Data and Code Availability

The raw data generated for this study is available from the corresponding author (MKJ) upon reasonable request. Derived datasets supporting the current findings and the custom codes used for data analysis are available on the github page: https://github.com/csbBSSE/Phenotypic_plasticity.

## Supplementary Information

### 1. Validating RACIPE simulation parameters

As mentioned in methods, we used the default simulation parameters except for the number of network parameter sets to be chosen. Among other simulation parameters, the integration method and the number of initial conditions at which each network parameter set is simulated have the potential to impact the phenotypic distributions significantly. Hence, to check the effects of these two simulation parameters, we chose networks representative of the network sizes considered in the paper, i.e, GRHL2 to represent small networks, NRF2 to represent medium sized networks and EMT RACIPE to represent the large networks. We then simulated each of these networks in 4 ways:

- RACIPE100: RACIPE with 10000 parameter sets and default values for all other simulation parameters (specifically, 100 initial conditions per parameter set and Euler’s method of integration).
- Matlab100: ode23s algorithm [**?**] in MATLAB R2018b (Mathworks) with 100 initial conditions and same parameter sets as obtained from RACIPE100 simulations. Since ode23s is a stiff system solver, this simulation gives us the effect of stiff systems the can potentially arise due to random sampling of network parameter sets.
- Matlab1000: Matlab ode23s with 1000 initial conditions and same parameter sets as above. This, in comparison with the previous simulations, reflects the effect of number of initial conditions used to obtain the steady states.
- RACIPE1000: RACIPE with 1000 simulations to check the effects of initial conditions in the operating domain of RACIPE. Since the RACIPE realization here is different from RACIPE100 simulations, the parameter sets need not be the same.

In Matlab simulations, the steady state was said to have reached when state change over 10 time-steps was less than 0.001. All the simulations were run in triplicates. We then compared the discretized phenotypic frequencies from these simulations in pairs, and measured:

- Fold change in the phenotype frequency levels using volcano plots, with p-values obtained from t-test on n=3 replicates. The null hypothesis is that the mean log fold change is 0.
- JSD of mean phenotypic distributions.

The results, shown in **Fig S7**, suggest that while these simulation parameters can affect frequencies of individual phenotypes, the changes at an ensemble level, as measured by JSD, are minute.

**Figure S1:**
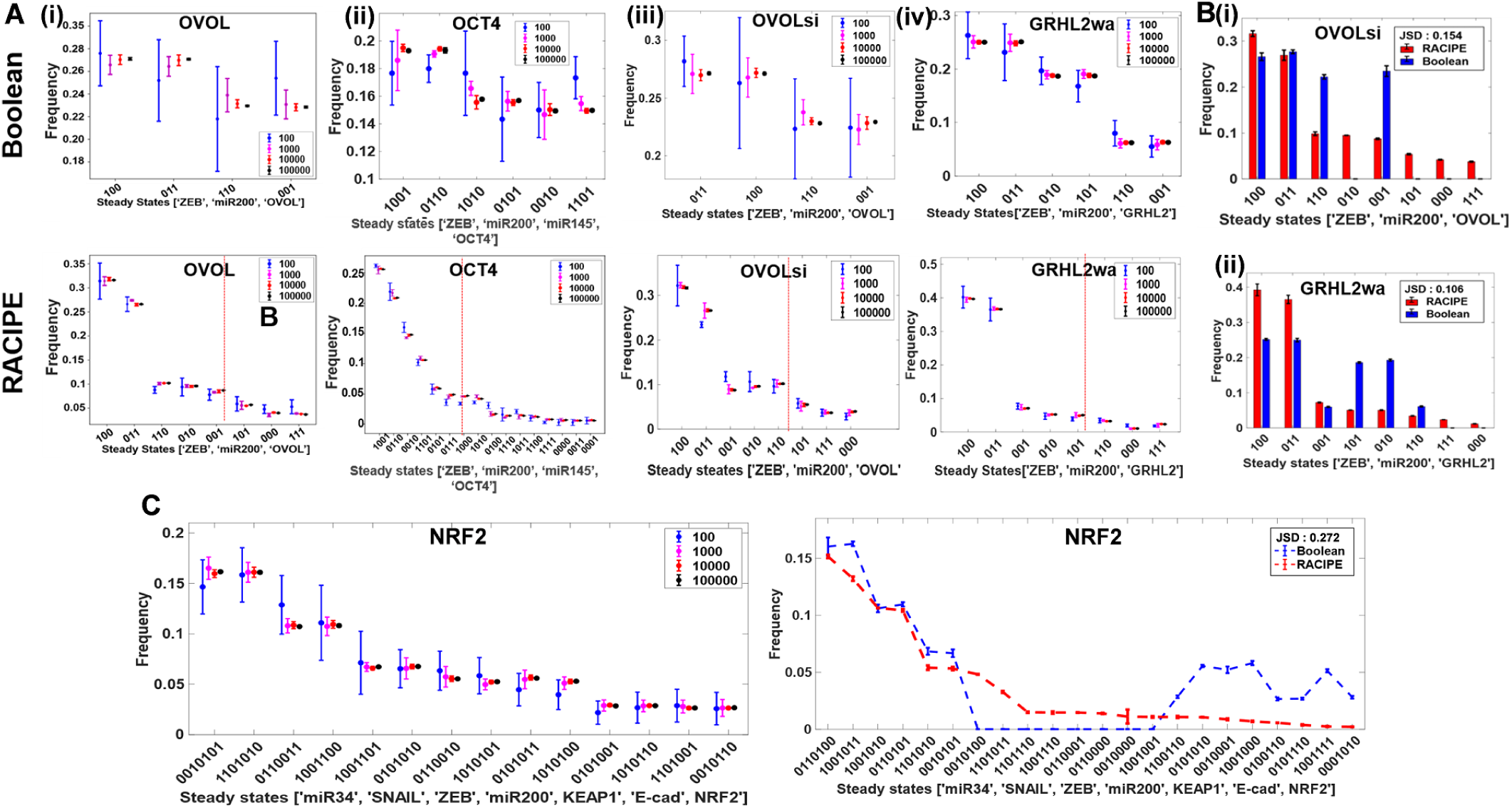
Quantitative convergence (QC) and comparison of phenotypic distributions obtained from RACIPE and Boolean A. QC of (i) OVOL, (ii) OCT4, (iii) OVOLsi and (iv) GRHL2wa using Boolean (top) and RACIPE (bottom). B. Comparision of steady state frequencies obtained from RACIPE and Boolean for GRHL2wa and OVOLsi C. NRF2 QC plot (Boolean) and comparison of RACIPE and Boolean state frequency distributions

**Figure S2:**
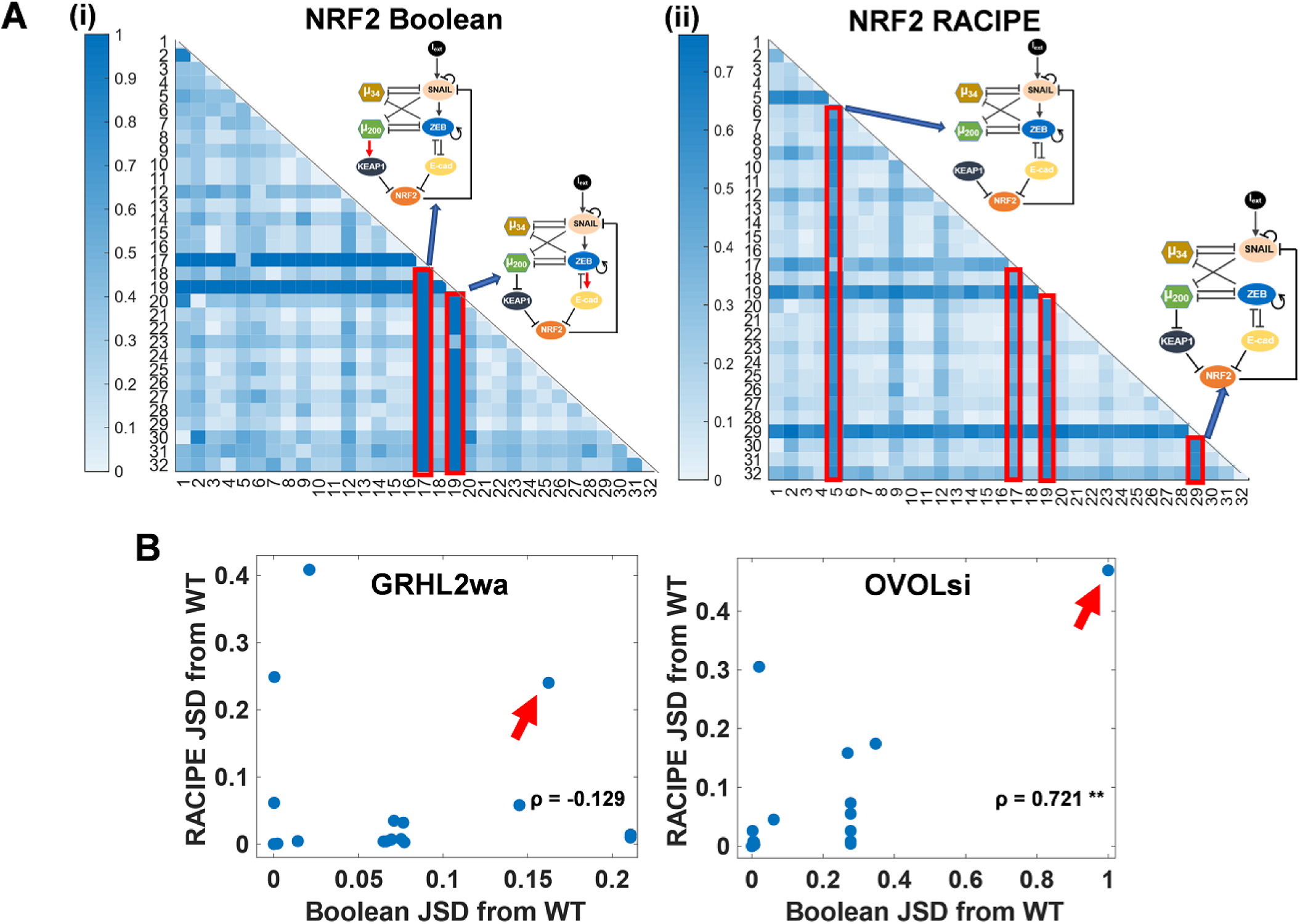
Effect of perturbations on phenotype distribution A. Heatmap of JSD of perturbations made to NRF2 network from each other (no edge additions) as obtained from i) Boolean and ii) RACIPE. The strongest perturbations are highlighted and the corresponding networks are shown with the heatmap. B. Scatter plots of JSD of perturbed networks from WT obtained from Boolean and RACIPE for GRHL2wa and OVOLsi.

**Figure S3:**
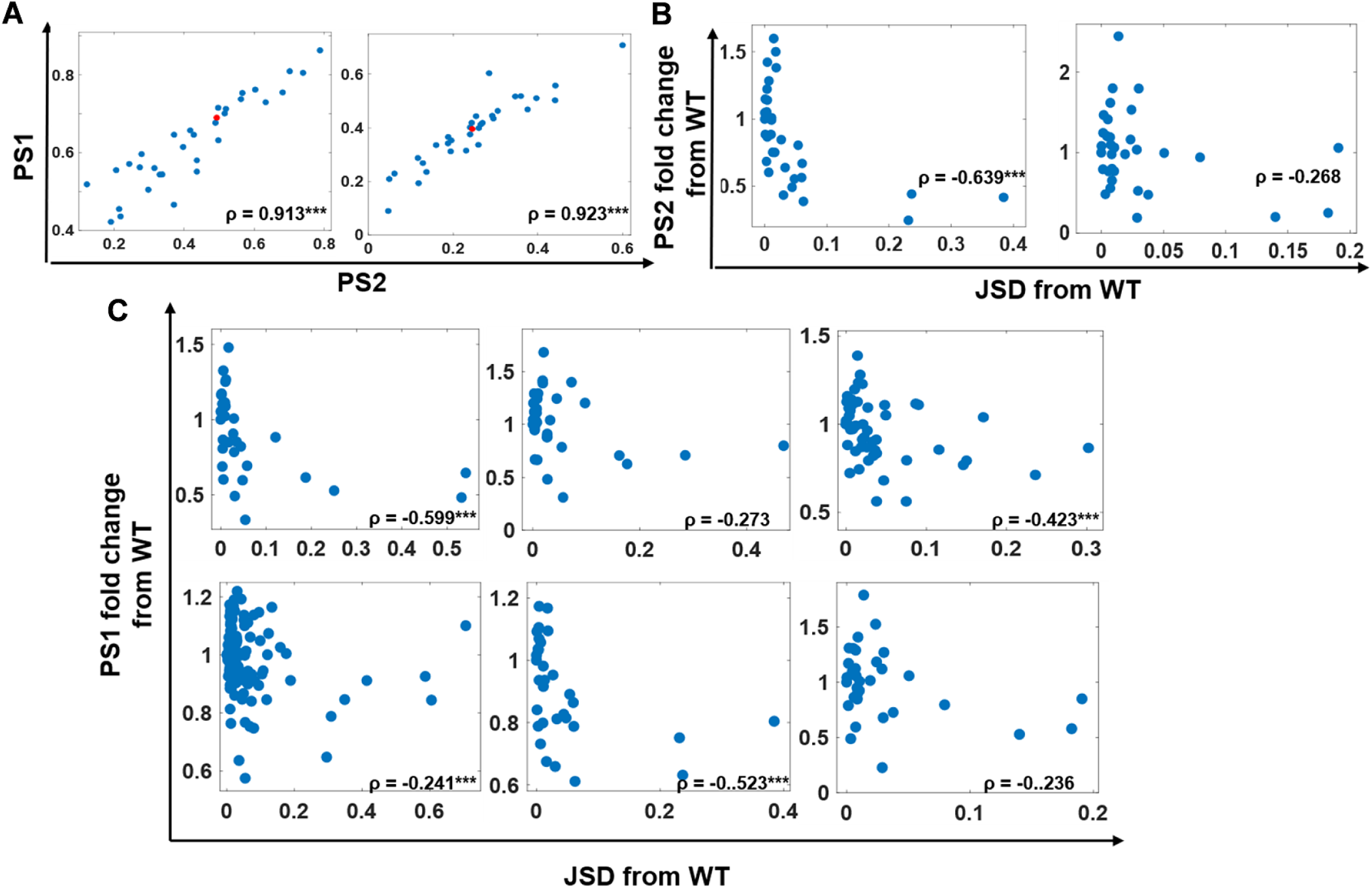
Effect of JSD on phenotypic plasticity A. Scatter plot between PS1 and PS2 for GRHL2wa and OVOLsi. B. Scatter plots for GRHL2wa and OVOLsi between fold change in plasticity vs change in phenotypic distribution (JSD). Each dot represents a perturbed network topology of the mentioned network. C. Same plots as B for all the networks, using PS1 instead of PS2

**Figure S4:**
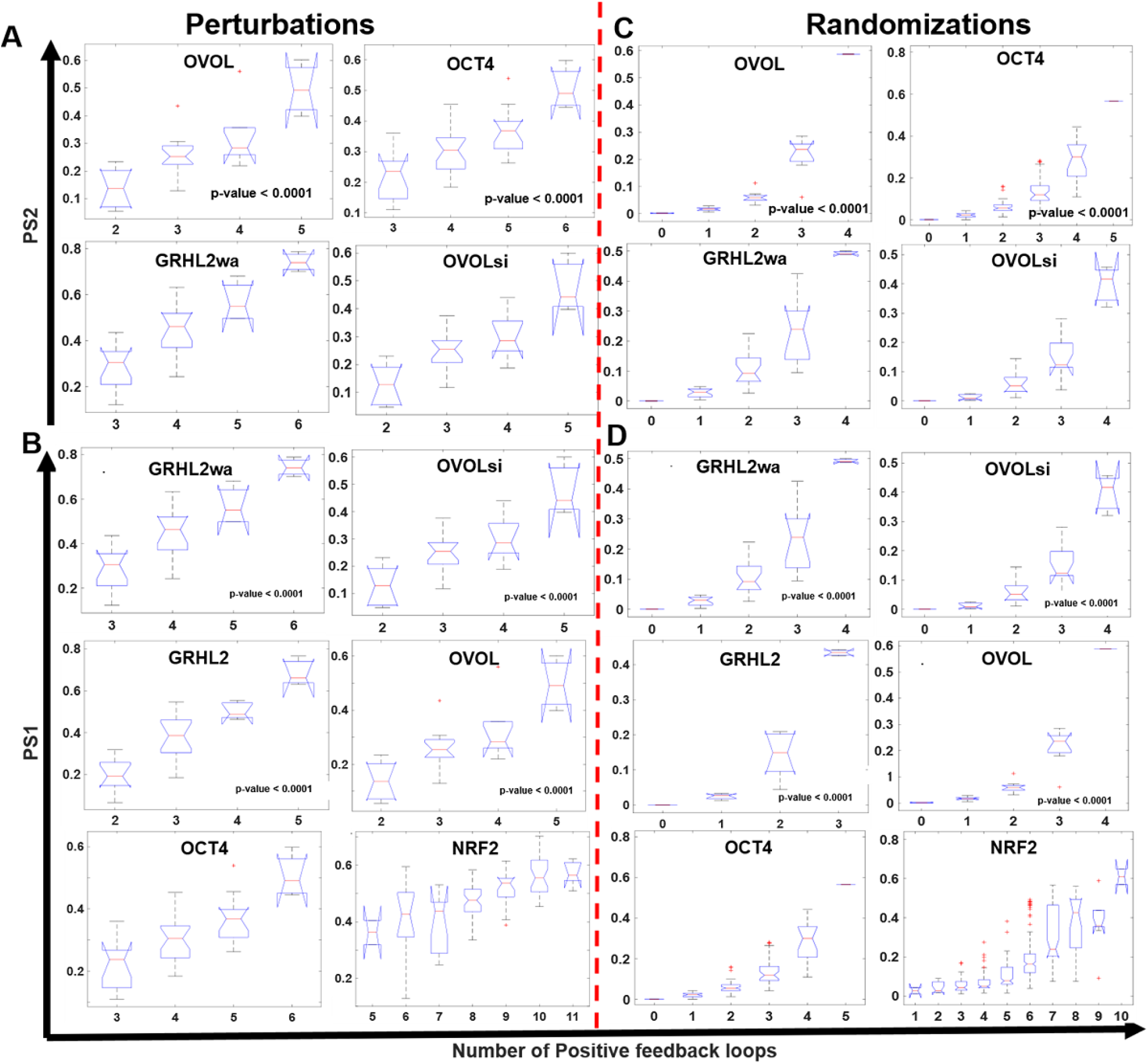
Effect of positive feedback loops on plasticity A. Box-plot of PS2 vs positive cycles for GRHL2wa and OVOLsi perturbed network topologies B. Box-plots of PS1 vs positive cycles for all perturbed network topologies coresponding to each EMP network C. Same as A but randomized circuits D. Same as B but randomized circuits

**Figure S5:**
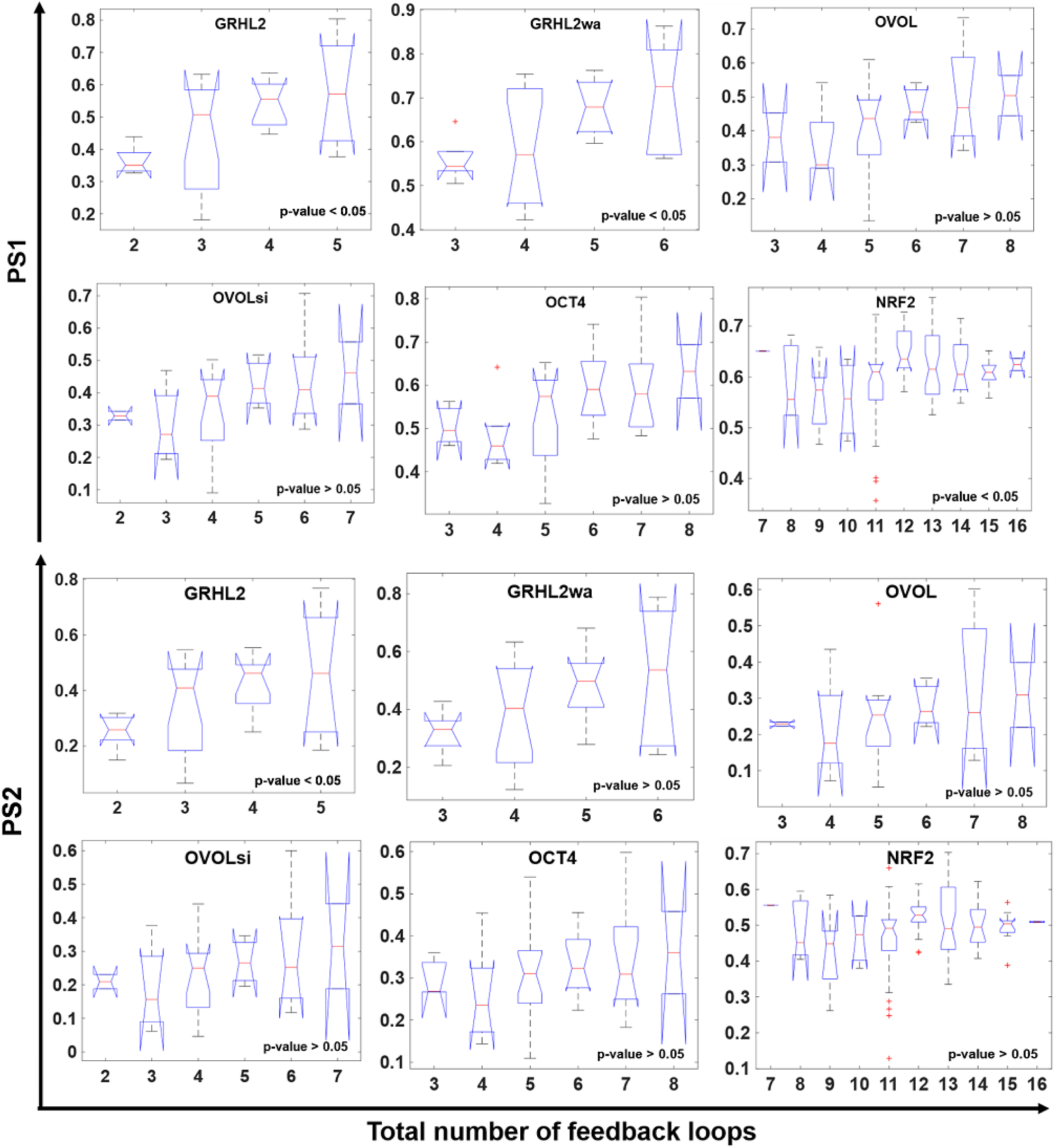
Effect of total number of feedback loops on network plasticity for perturbations of all 6 networks.

**Figure S6:**
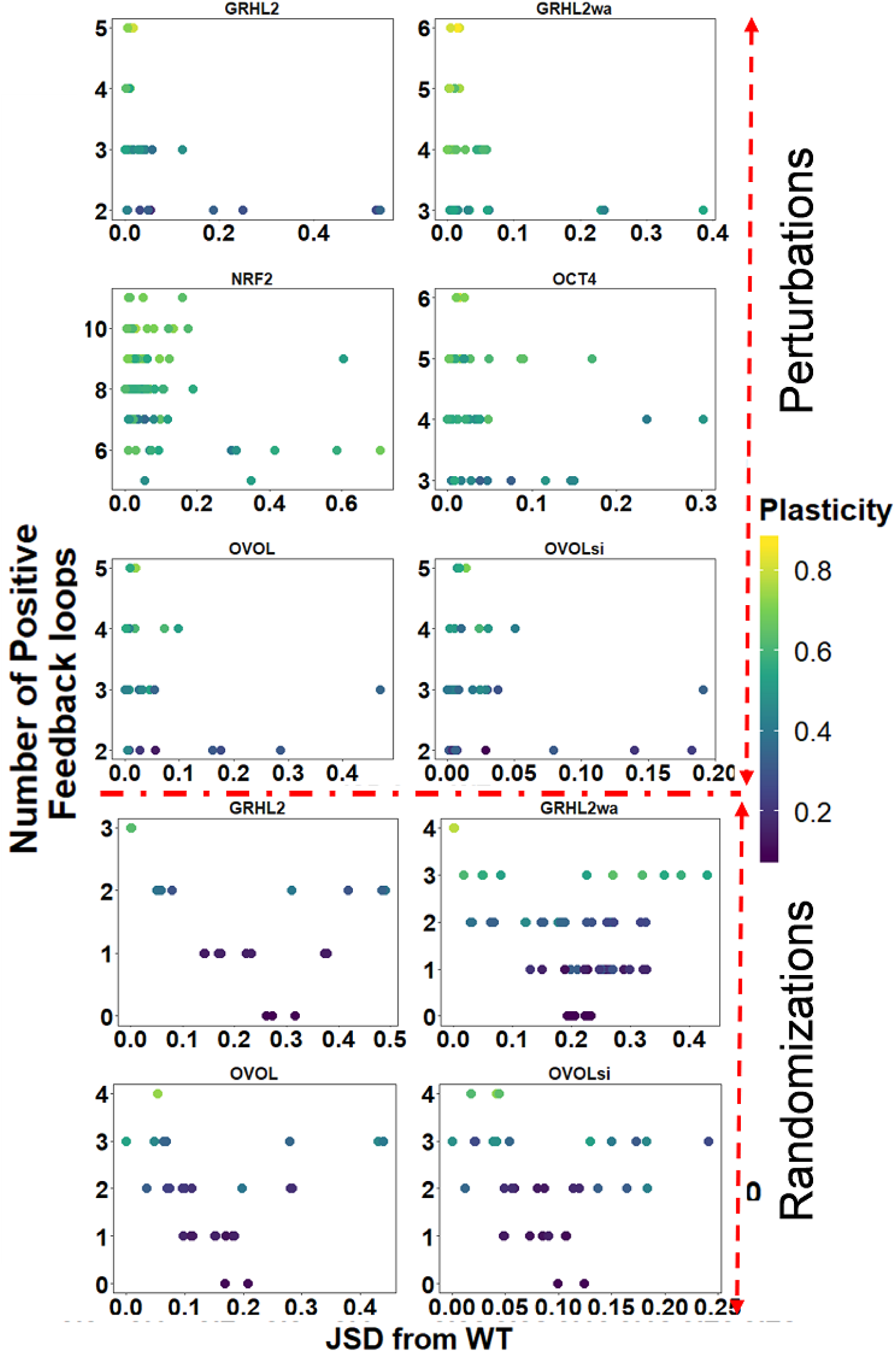
Combined effect of positive cycles and JSD on plasticity A. Positive cycles vs JSD scatter plots for perturbed network topologies of all 6 networks, colored according to plasticity B. Same as A but for randomized circuits

**Figure S7:**
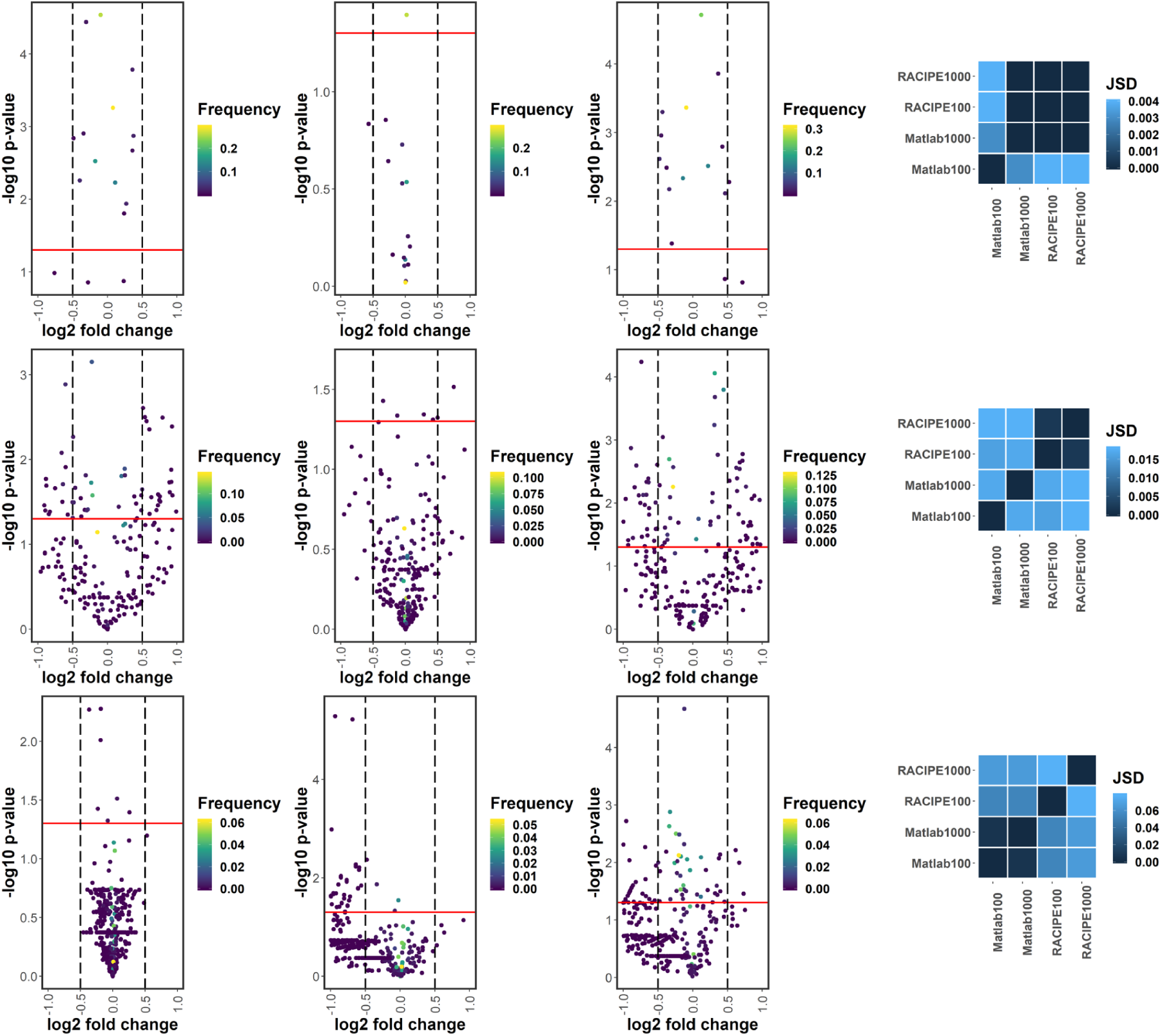
Testing RACIPE simualtion parameters. A. Results for GRHL2. Volcano plots compare fold changes in steady states from (i) Matlab100 and MATLAB1000, (ii) RACIPE100 and RACIPE 1000, (iii)RACIPE100 and MATLAB100. Each point represents a steady state. The red horizontal line corresponds to p=0.05. Fold change in any steady state below the red line is insignificant. The vertical lines correspond to a fold change of 0.7 and 1.41 respectively. Each steady state has been colored according its observed frequency. (iv) JSD heatmap of the mean phenotypic distributions obtained from all 4 simulations. B. Same as A but for NRF2. C. Same as A but for EMT_RACIPE

**Table S1:** GRHL2 single-edge perturbations including edge-additions. A given single edge perturbation is named as: FromNode-ToNode_OriginalEdge-PerturbedEdge. 0: no edge, 1: activation, 2: inhibition. Network number 2 is ‘wild-type’

| Label | Network |
| --- | --- |
| 1 | GRHL2-GRHL2_0-1 |
| 3 | ZEB-miR200_2-1 |
| 4 | GRHL2-GRHL2_0-2 |
| 5 | miR200-miR200_0-1 |
| 6 | miR200-ZEB_2-0 |
| 7 | miR200-ZEB_2-1 |
| 8 | SNAIL-SNAIL_0-1 |
| 9 | ZEB-miR200_2-0 |
| 10 | ZEB-ZEB_1-0 |
| 11 | ZEB-ZEB_1-2 |
| 12 | ZEB-GRHL2_2-0 |
| 13 | ZEB-GRHL2_2-1 |
| 14 | SNAIL-GRHL2_0-1 |
| 15 | SNAIL-GRHL2_0-2 |
| 16 | GRHL2-ZEB_2-0 |
| 17 | GRHL2-ZEB_2-1 |
| 18 | SNAIL-ZEB_1-2 |
| 19 | GRHL2-miR200_0-1 |
| 20 | GRHL2-SNAIL_0-2 |
| 21 | GRHL2-miR200_0-2 |
| 22 | GRHL2-SNAIL_0-1 |
| 23 | SNAIL-ZEB_0-1 |
| 24 | miR200-GRHL2_0-1 |
| 25 | SNAIL-ZEB_0-2 |
| 26 | SNAIL-ZEB_1-0 |
| 27 | miR200-GRHL2_0-2 |
| 28 | miR200-miR200_0-2 |
| 29 | miR200-SNAIL_0-1 |
| 30 | miR200-SNAIL_0-2 |
| 31 | SNAIL-miR200_2-0 |
| 32 | SNAIL-miR200_2-1 |

**Table S2:**
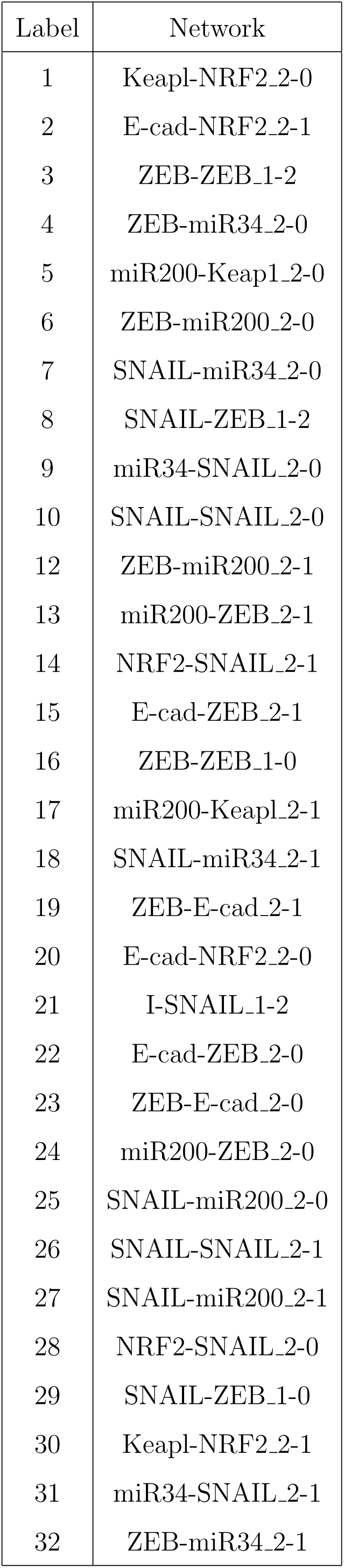
NRF2 perturbations without edge-additions. A given single edge perturbation is named as: FromNode-ToNode_PreviousEdge-OriginalEdge. 0: no edge, 1: activation, 2: inhibition. Network number 11 is ‘wild-type’

| Label | Network |
| --- | --- |
| 1 | Keap1-NRF2_2-0 |
| 2 | E-cad-NRF2_2-1 |
| 3 | ZEB-ZEB_1-2 |
| 4 | ZEB-miR34_2-0 |
| 5 | miR200-Keap1_2-0 |
| 6 | ZEB-miR200_2-0 |
| 7 | SNAIL-miR34_2-0 |
| 8 | SNAIL-ZEB_1-2 |
| 9 | miR34-SNAIL_2-0 |
| 10 | SNAIL-SNAIL_2-0 |
| 12 | ZEB-miR200_2-1 |
| 13 | miR200-ZEB_2-1 |
| 14 | NRF2-SNAIL_2-1 |
| 15 | E-cad-ZEB_2-1 |
| 16 | ZEB-ZEB_1-0 |
| 17 | miR200-Keap1_2-1 |
| 18 | SNAIL-miR34_2-1 |
| 19 | ZEB-E-cad_2-1 |
| 20 | E-cad-NRF2_2-0 |
| 21 | I-SNAIL_1-2 |
| 22 | E-cad-ZEB_2-0 |
| 23 | ZEB-E-cad_2-0 |
| 24 | miR200-ZEB_2-0 |
| 25 | SNAIL-miR200_2-0 |
| 26 | SNAIL-SNAIL_2-1 |
| 27 | SNAIL-miR200_2-1 |
| 28 | NRF2-SNAIL_2-0 |
| 29 | SNAIL-ZEB_1-0 |
| 30 | Keap1-NRF2_2-1 |
| 31 | miR34-SNAIL_2-1 |
| 32 | ZEB-miR34_2-1 |

